# Belowground Herbivory Shapes Carbon Release from Tree Roots and Its Retention in Soil

**DOI:** 10.64898/2026.11.25.768934

**Authors:** Jaeyoung Yoo, Taryn L. Bauerle, Richard P. Phillips, André L.C. Franco

## Abstract

Belowground root inputs contribute to stable soil organic carbon (SOC) significantly more than aboveground plant inputs. Yet, current understanding of the role of belowground root herbivory in ecosystem C cycling is remarkably weaker than that of aboveground herbivory and litter inputs. Here, we experimentally exposed roots of co-occurring mature hickory (*Carya*) and maple (*Acer*) trees, in a temperate hardwood forest, to increasing herbivory pressure by ubiquitous root-feeding soil nematodes. We investigated responses and underlying mechanisms of root C exudation as well as downstream indirect effects of root herbivory on mineral-associated organic carbon (MAOC) pools. Increased root herbivory promoted C exudation, which was strongly associated with increasing root necrosis and resulted in indirect positive effects on MAOC. Importantly, these direct and indirect effects were larger in magnitude and only significant in hickory but not maple. Instead, the latter appeared to channel herbivory response toward diversified chemical defense signaling rather than bulk exudate release, while hickory’s more extensive root damage and increased exudation rate with less compositional change suggested a greater role of passive exudate release through “leaking wounds” in roots. Taken together, these findings indicate divergent eco-physiological mechanisms and ecosystem C consequences of root herbivory in mature hickory and maple trees, and show that changes in exudate quantity rather than chemical composition impact soil C. This study extends field-based causal explanation of root C inputs and retention in soil by highlighting the role of root-feeding animals in regulating rhizodeposition, and offers a basis for integrating belowground herbivory into multitrophic biogeochemistry.

**Significance Statement:** Root-derived carbon is a major source of persistent soil carbon, yet the role of root-feeding animals in regulating this carbon pathway remains overlooked. We show experimentally in a temperate forest that nematodes feeding on the absorptive tree roots alter fine-root carbon release into soil, with consequences for persistent soil carbon. Co-occurring hickory and maple trees responded through contrasting mechanisms: herbivory promoted root damage and carbon leakage in hickory, leading to an increase in the persistent soil carbon pool, while maple responded primarily by increasing defense-related compounds without changes in exudation rate and soil carbon. These findings identify belowground herbivory as an underappreciated pathway linking plant-soil interactions to soil carbon persistence and suggest that tree identity determines how this pathway operates.

## Introduction

Research suggesting herbivory as a major driver of soil C focuses largely on aboveground herbivory increasing plant inputs to soil (1, 2). However, given that root-feeding animals such as nematodes can consume more biomass than aboveground grazers (3), and root C inputs are up to five times more likely than aboveground inputs to be incorporated into persistent SOM (4), more studies of how root herbivory affects soil C are needed. An estimated 46% of root exudates can be transformed into persistent SOC (decadal–millennial timescales), thus contributing to long-term C sequestration in soil, compared with only 9% of root tissues and 7% of aboveground residues (5). Thus, understanding how belowground ecological interactions regulate rhizodeposition has increasingly gained attention (6).

The role of belowground herbivores in C exudation and retention in soil remains remarkably understudied. Existing evidence suggests that root herbivory by nematodes can enhance the allocation of recently photoassimilated C to roots (7), thereby increasing root exudation and stimulating rhizosphere microbial activity (8, 9). This suggests that roots that are affected by root feeding damage may increase cellular leakage of root-derived C into soil, potentially altering SOC formation and persistence pathways. Therefore, root herbivory by nematodes may be an overlooked regulator of belowground C inputs and retention through rhizodeposits, and a missing component in conceptual and predictive ecosystem C frameworks.

Root exudation accounts for 7–14% of global annual gross primary productivity (10) and plays a significant role in shaping SOC formation and persistence (11, 12). Exudates alter the formation, stabilization and turnover of MAOC – the fraction of C considered to be more persistent (13). Slight to moderate increases in C exudation generally promote MAOC accumulation, while high exudation rates can accelerate MAOC turnover, elevating both assimilation and loss with little MAOC net change (14, 15). Exudate composition could also alter MAOC dynamics. Sugars, for example, can increase microbial assimilation and priming of MAOC (14, 16), whereas organic acids and amino acids can alter rhizosphere microbial communities, promoting MAOC accumulation (14, 16). Phenolic compounds can also alter soil microbial communities, with direct consequences for MAOC formation and persistence (17, 18). Thus, belowground ecological interactions that modify root exudation rates or composition could have downstream effects on SOC accrual and persistence.

Emerging evidence points to root-feeding soil animals as potential regulators of both the rate (8) and chemical composition (19) of root exudation. Studies in herbaceous species have found that root-feeding nematodes can damage root structure and morphology, creating “leaking wounds” that passively increase exudation (8, 9). Plants can also selectively exude metabolites as an active defense mechanism against herbivore infection, increasing antimicrobial or antiherbivore compounds and attractants for beneficial rhizobacteria (17). For example, barley has been reported to increase sugar concentrations in root exudates following infection by root-knot nematodes (*Meloidogyne incognita*) (19). These pieces of evidence suggest that root herbivory may represent a neglected pathway controlling root exudation. With anthropogenic changes in climate (20), nutrient pollution (21), and management (22) resulting in increased root-feeding nematode populations and outbreaks, plants in several types of ecosystems are increasingly exposed to greater belowground herbivory. Understanding how root-feeding animals regulate rhizosphere C fluxes through mechanisms such as passive exudate leakage and active plant defense across dominant plant taxa can improve predictions of soil C storage and ecosystem persistence.

Research on nematode herbivory has historically focused on pest control and yield in croplands, where nematodes cause over 12% annual crop yield losses worldwide, and single-species interactions are easier to isolate than in complex, high biodiversity ecosystems such as forests (23). Root herbivory has received less attention in forests, partly because complex soil food webs are assumed to dilute herbivore impacts, energy flow is considered dominated by bacterial and fungal decomposition channels rather than direct root-herbivore channels, and effects on long-lived trees are presumed negligible relative to annual crops. However, root-feeding nematodes often dominate temperate forest soil fauna communities and are particularly abundant under tree species that associate with arbuscular mycorrhizal (AM) fungi, such as *Acer saccharum* (24). Whether nematodes affect tree species that associate with the other dominant type of mycorrhizal fungi in forests – ectomycorrhizal (ECM) fungi – is unknown. Unlike AM fungi, which penetrate root cells, ECM fungi form a hyphal mantle around absorptive roots. As such, ECM fungi may physically block the release of exudates into soil from reaching the soil(25) and impedes endoparasitic nematodes from locating and penetrating the root epidermis with their feeding stylets. Therefore, root-feeding nematodes may have an impactful role in regulating tree C inputs into soil through exudation in forest ecosystems, with the magnitude of their influence potentially mediated by tree type.

Here, we experimentally tested whether increasing belowground herbivory in a temperate hardwood forest causes directional and taxon-specific changes in root C exudation rates and chemical composition of mature trees and whether those changes have consequences on soil C pools. Under field conditions, we exposed roots of two dominant tree types, hickory (ECM-associated) and maple (AM-associated), to increasing populations of root-feeding nematodes to investigate the magnitude and mechanisms behind root exudation responses. We asked: 1) What are the effects and underlying mechanisms of root herbivory on tree exudation? 2) Do co-occurring hickory and maple trees differ in their responses to root herbivory? 3) Does root herbivory indirectly alter persistent soil C pools?

## Results

### Root herbivory effects on root morphology

The proportion of necrotic points in roots increased with herbivory pressure, but that effect was marginally dependent on tree type; necrosis rate increased with herbivory in hickory but not maple (herbivory × tree type: p = 0.089; marginal R^2^ = 0.081; conditional R^2^ = 0.139; Fig. S1). Root necrosis rate was also positively associated with C exudation rate (y = 0.51x – 3.24, R^2^ = 0.219, p = 0.002), with a stronger relationship for hickory (y = 1.05x – 18.01, R^2^ = 0.493, *p* = 0.001) than maple (y = 0.23x + 3.78, R^2^ = 0.074, *p* = 0.199; Fig. 1). Other root morphological traits including mean root diameter, specific root length, branching frequency, and number of root tips were not significantly affected by herbivory (Fig. S2).

**Fig. 1.**
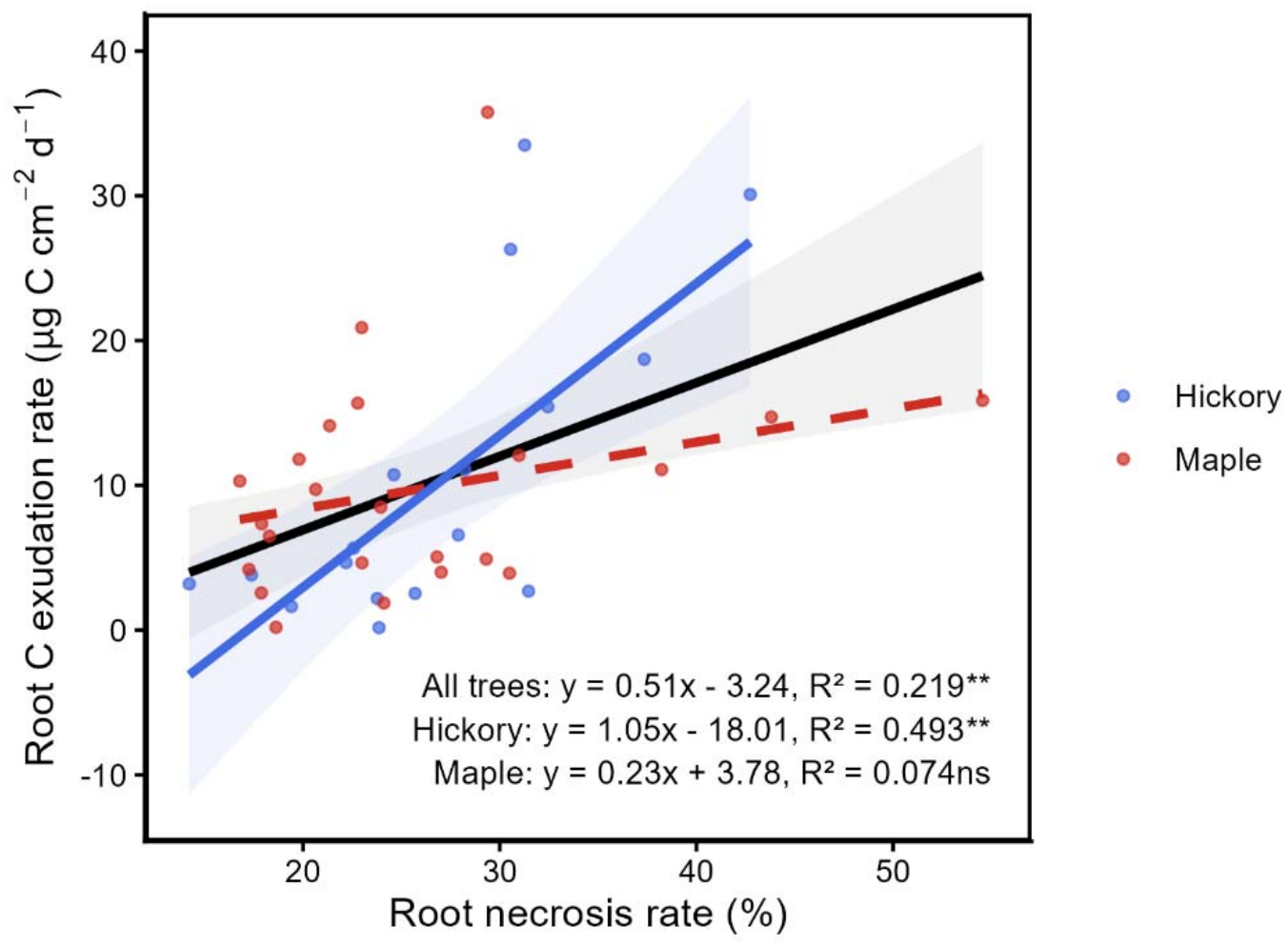
Changes in root C exudation rate by root necrosis rate in hickory and maple plots (n=54). Regression lines indicate the relationship between root necrosis rate and root C exudation rate, with solid lines representing significant effects (p□<□0.05) and dashed lines indicating non-significant trends (p□≥□0.05). The black line indicates the regression line for all data included, the blue line for hickory plot data, and the red line for maple plot data.

### Root herbivory effects on root C exudation rate and metabolomes

The effect of increasing root herbivory pressure on C exudation rate varied by tree type (herbivory × tree type: p = 0.023; marginal R^2^ = 0.232; conditional R^2^ = 0.352). Root C exudation increased linearly with increasing root herbivory pressure in hickory (y = 0.013x + 0.946), whereas maple showed little response (y = −0.003x + 9.508) (Fig. 2).

**Fig. 2.**
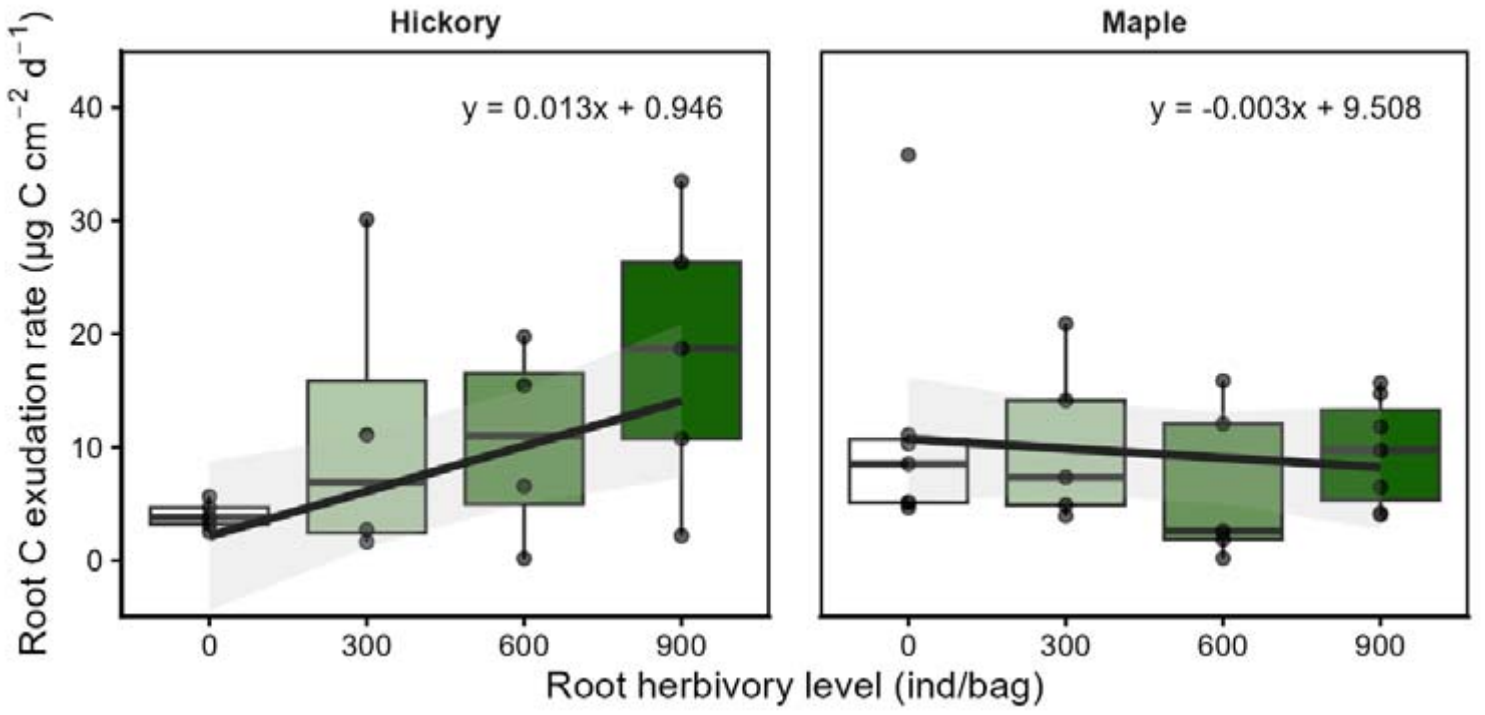
Root C exudation rates (µg C cm^−2^ d^−1^) across root herbivory levels (0, 300, 600, and 900 nematodes per microcosm) in hickory and maple plots (n=42). Boxes represent the interquartile range (IQR), with the center line indicating the median. Whiskers extend to 1.5 × IQR, and individual points represent biological replicates. Darker green shading indicates higher herbivory levels. The solid line represents a linear regression fit with 95% confidence intervals (shaded area).

Changes in exudation composition in response to herbivory also differed between the two species. We targeted negative ionization mode root metabolome (26), as negative mode ionization has higher efficiency in capturing compounds that deprotonate, such as sugars, organic acids, and phenolic compounds (27, 28) that are relevant to root defense responses; we recognize that positive mode ionization can also contain compounds that respond to herbivory and therefore present these compounds as supplementary information. Although exudate compound richness was unaffected by herbivory level (Fig. S3), exudate composition (based on negative ionization mode; see methods) varied with root herbivory pressure and tree type (PERMANOVA; herbivory × tree type: F = 1.841, p = 0.045, R^2^ = 0.045; Fig. 3A). Herbivory explained substantially more compositional variation (F = 17.611, p < 0.001, R^2^ = 0.453) than tree type (F = 11.106, p < 0.001, R^2^ = 0.095). Phenolics dominated exudates across all herbivory levels and had the greatest increase under high herbivory pressure compared with other metabolites (Fig. 3B). These patterns were generally consistent in positive ionization mode (Figs. S4 and S5). Of the four metabolite groups known to regulate soil C dynamics, concentrations of sugars, organic acids, and phenolics increased non-linearly with herbivory, whereas amino acid concentrations did not change (sugar: F = 2.48, p = 0.094, marginal R^2^ = 0.338; organic acid: F = 4.30, p = 0.019, marginal R^2^ = 0.283; amino acid: F = 1.60, p = 0.211, marginal R^2^ = 0.290; phenolics: F = 18.86, p < 0.001, marginal R^2^ = 0.505; Figs. S6). There was no effect of tree types or interaction between herbivory level and tree type. These patterns were generally consistent in positive ionization mode (Fig. S7).

**Fig. 3.**
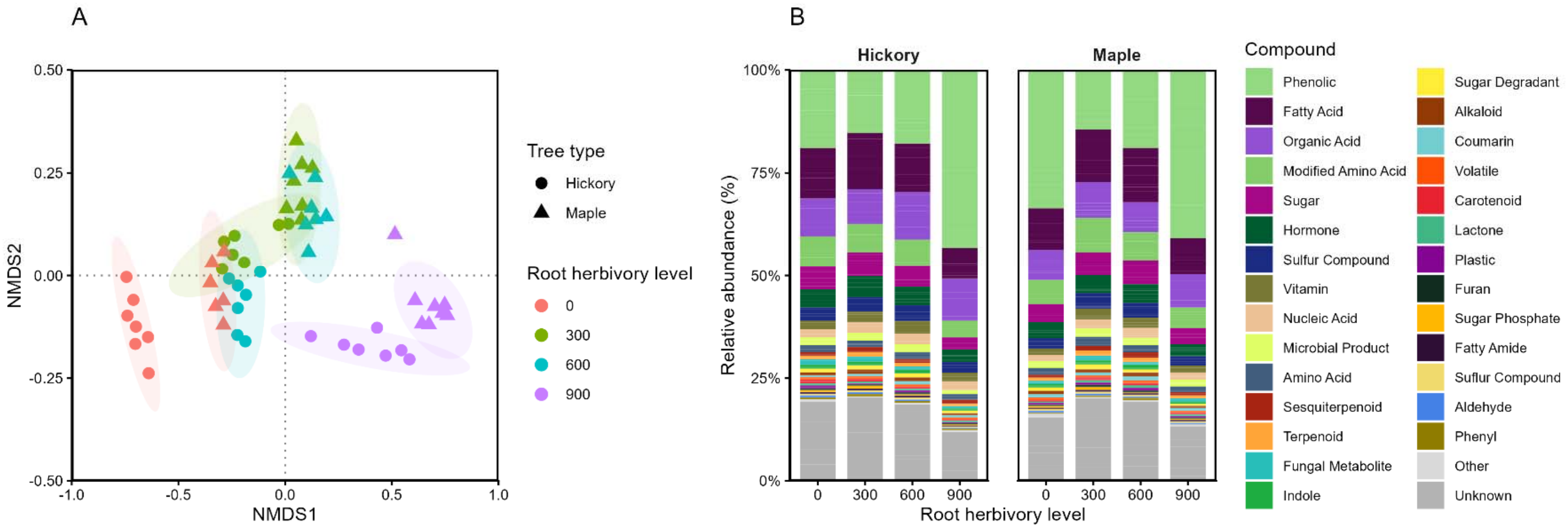
Non-metric multidimensional scaling (NMDS) ordination of root exudate composition (A) and relative abundance (%) of root exudate compound classes (B) in hickory and maple plots across root herbivory levels (0, 300, 600, and 900 nematodes per microcosm) in negative ionization mode (n=56). (B) Each point in NMDS ordination plot represents one biological replicate. Symbols denote tree type (circle = hickory; triangle = maple) and colors indicate herbivory level (red = 0, green = 300, teal = 600, purple = 900 nematodes per microcosm). Ellipses represent 95% confidence intervals around group centroids by herbivory level. (B) Each bar in relative abundance plot represents the mean relative abundance of identified compound classes per herbivory level. Colors denote compound classes as indicated in the legend. Gray bars represent compounds of unknown classification.

Indicator species analysis identified one and five indicator compounds for hickory and maple in negative mode, respectively (Table S1; Figs. S8–S11). Those included vitamin A derivative in hickory (IndVal = 0.908, p = 0.031), and benzoic acid (IndVal = 0.990, p = 0.029), vanilpyruvic acid (IndVal = 0.929, p = 0.038), nicotinic acid (IndVal = 0.948, p = 0.006), 6-hydroxynicotinic acid (IndVal = 0.940, p = 0.003) and 9,10-dihydroxystearic acid in maple (IndVal = 0.949, p = 0.005; Table S1). Indicator species for hickory and maple in positive mode are listed in Table S2.

### Downstream effects on soil C

The effects of root herbivory on SOM fractions and stoichiometry differed between tree types (Fig. S12). Neither MAOC nor particulate organic C (POC) responded directly to root herbivory treatments (MAOC: F = 0.014, p = 0.907; POC: F = 0.099, p = 0.754), and neither showed a herbivory × tree type interaction (MAOC: F = 0.531, p = 0.470; POC: F = 0.454, p = 0.504). However, POC was higher in hickory than in maple soils (F = 21.41, p < 0.001). In contrast, C:N ratios of both SOM fractions showed significant herbivory × tree type interactions. C:N ratio of mineral-associated organic matter (MAOM) increased with herbivory pressure in hickory soils but declined in maple soils (herbivory × tree type interaction: F = 18.09, p < 0.001, marginal R^2^ = 0.157, conditional R^2^ = 0.502). The C:N ratio of particulate organic matter (POM) showed a similar pattern but with shallower slopes (herbivory × tree type interaction: F = 8.56, p = 0.005, marginal R^2^ = 0.162, conditional R^2^ = 0.468).

Structural equation models (SEMs) showed acceptable fit for hickory (Fisher’s C = 16.778, p = 0.079) and maple (Fisher’s C = 14.348, p = 0.158), indicating that the hypothesized indirect pathway by which root herbivory mediates MAOC were consistent with the observed data (Fig. 4). In hickory, root herbivory significantly predicted root exudate composition (β = 0.833, p < 0.001) and showed a marginally significant positive effect on root necrosis rate (β = 0.340, p = 0.089). Greater root necrosis rate predicted higher root C exudation rate (β = 0.659, p = 0.001), which in turn predicted greater MAOC (β = 0.546, p = 0.004). By contrast, exudate composition did not predict root C exudation rate in hickory (β = 0.058, p = 0.745). Thus, despite no direct herbivory treatment effects on MAOC (Fig. S12B), we found support for an indirect pathway linking herbivory to MAOC through root necrosis and increased root C exudation in hickory trees. In maple, root herbivory predicted root exudate composition (β = 0.970, p < 0.001), but not root necrosis rate (β = 0.005, p = 0.793). Neither root necrosis rate (β = 0.037, p = 0.827) nor root exudate composition (β = −0.076, p = 0.725) predicted root C exudation rate, and root C exudation rate did not predict MAOC (β = −0.123, p = 0.538). Thus, unlike hickory, the maple SEM provided no evidence of indirect root herbivory controls on MAOC through root necrosis rate or root C exudation rate.

**Fig. 4.**
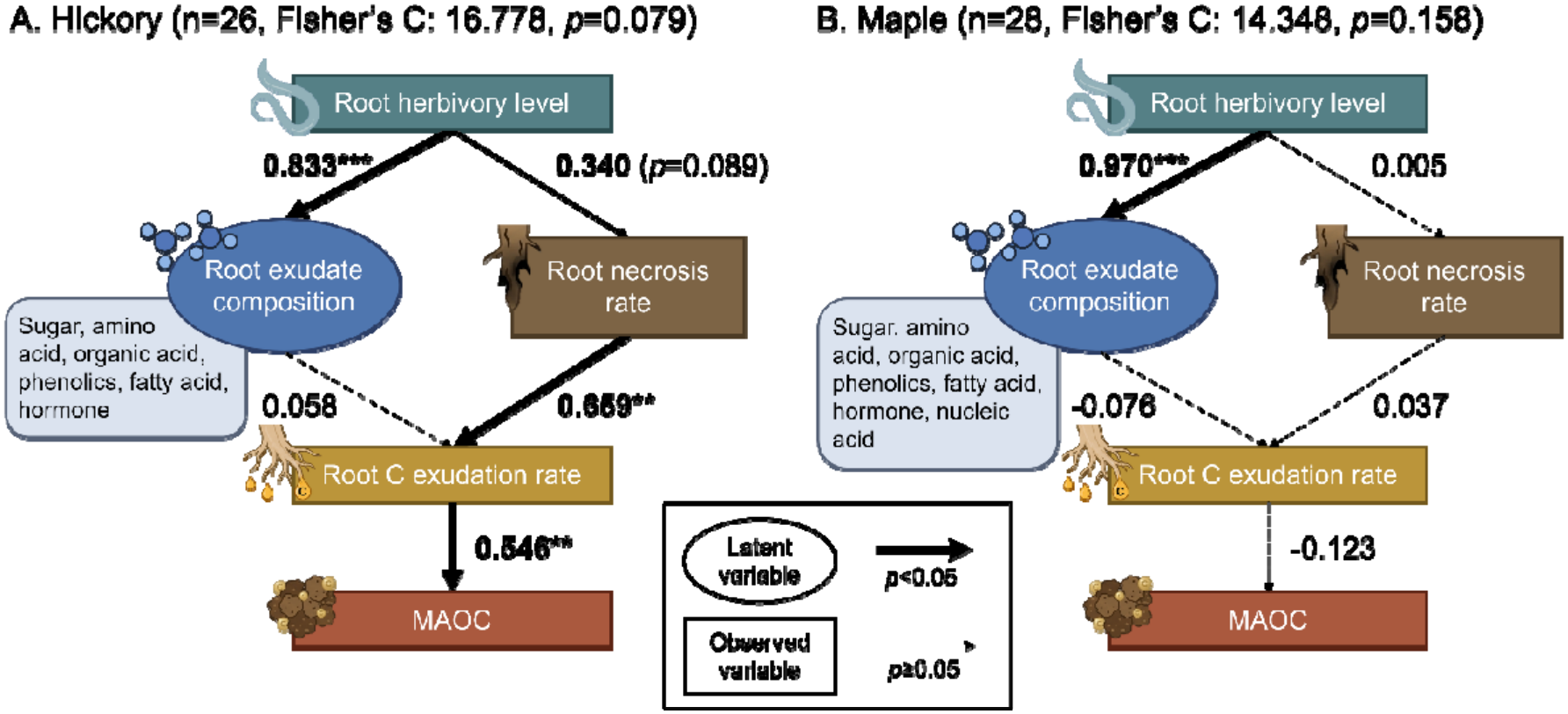
Piecewise structural equation models (SEMs) linking root herbivory to MAOC via root exudate composition, root necrosis rate, and root C exudation rate in (A) hickory and (B) maple. Ellipses represent latent variables, and rectangles represent observed variables. Solid arrows indicate statistically significant paths (p < 0.05) and dashed arrows indicate non-significant paths (p ≥ 0.05) except root herbivory level ~ root necrosis rate in (A) hickory is presented as a thinner solid arrow since it has a marginally significant relationship (p = 0.089); arrow width is proportional to the magnitude of the standardized path coefficient (β), shown alongside each path. The light blue boxes list the metabolite classes comprising the root exudate composition latent variable for each tree species. Significance codes: ** p < 0.01, *** p < 0.001.

## Discussion

### Taxon-specific responses to root herbivory: passive leakage in hickory versus active defense in maple roots

Root herbivory pressure drove tree genus-specific responses in root C exudation rate, metabolite composition, and root necrosis (Figs. 1, 2 and S1), suggesting different eco-physiological mechanisms by which hickory and maple trees respond to increased root herbivory. The different responses in hickory vs. maple may relate, in part, to their root traits. Hickory absorptive roots are characterized by thick diameter, high lignification, and slow root turnover (29, 30). Although thicker and more lignified roots could in theory provide greater physical resistance to herbivory, our results did not support this expectation. Instead, slower root turnover apparently constrains the capacity to replace or repair damaged fine-root tissues (31), consistent with the concurrent increases in root necrosis rate and C exudation rate observed under increasing herbivory (Figs. 1, 2 and S1). Our results also suggest that root-feeding nematodes may overcome the ECM fungal mantle surrounding hickory fine roots, potentially creating wounds through which exudates otherwise restricted by the mantle (25) are leaked. Conversely, maple roots are thinner and have higher root nitrogen and turnover rates (29, 30). Faster tissue replacement and repair (31) may explain the absence of detectable changes in maple root necrosis and C exudation rate found here (Figs. 1, 2 and S1). Thus, differences in root turnover emerge as a primary factor explaining the differential responsiveness of hickory and maple root morphology and exudation patterns to increased herbivory.

While our results suggest herbivory induced the passive release of exudates (i.e., through “leaking wounds”) in hickory, maple exudation rates were unaffected. However, exudate metabolite composition changed in maple without changes in total C exudation or root necrosis. Phenolic compounds became increasingly dominant in root exudates of both tree types under high herbivory pressure (Fig. 3B), consistent with a shift toward compounds potentially involved in antiherbivore and antimicrobial defense, microbial recruitment, allelopathic interactions, and nutrient acquisition (17, 32). However, maple trees exhibited a broader suite of defense-related compounds than hickory (Table S1 and S2). Hickory responses were concentrated in a single compound, vitamin A derivative, associated with signaling against abiotic and biotic stress generally, while maple responses involved more diverse defense pathways, including salicylic acid (i.e. benzoic acid), nicotinic acid (i.e. nicotinic acid and 6-hydroxynicotinic acid) signaling, as well as other putatively defense-related metabolites including 9,10-dihydroxystearic acid, and vanilpyruvic acid (Table S1), Positive mode data likewise showed a more diverse defense response in maple (Table S2). These patterns support the interpretation that maple channels its herbivory response towards diversified chemical signaling and active defense mechanisms rather than bulk exudate release, while hickory’s more extensive root damage leads to passive exudate release.

### Indirect effects of root herbivory on soil C are mediated by root damage and exudation rate

Our results indicate that, despite no detectable direct effects (Fig. S12A), root herbivory can indirectly affect MAOC through its effects on root morphology and C exudation rate (Fig. 4). Root exudates can both promote MAOC formation through direct mineral sorption and microbial processing, and destabilize existing MAOC through priming (15). Part of the existing evidence for net MAOC losses from exudate-induced priming comes from experiments of 30 days or less (14, 15), suggesting that (i) exudate-induced changes in MAOC can be detectable within a season, and that (ii) our 8 week-long experiment may have allowed sufficient time for offsetting initial MAOC priming by exudate C sorption and MAOC formation. Importantly, the pathway connecting root herbivory to MAOC through root necrosis and exudation rate was supported in hickory but not maple, suggesting that the increased passive leakage of root exudates observed in hickory was of sufficient magnitude to measurably increase MAOC concentration in soil. On the contrary, the active mobilization of defense metabolites without increased C exudation in maple did not translate into direct or indirect MAOC changes. These results indicate that the ecological impact of root herbivory on soil C is primarily influenced by effects on exudate quantity and not metabolite composition.

Changes in MAOM stoichiometry provide additional information on how herbivory altered MAOC accumulation. Increasing herbivory led to a higher C:N ratio of MAOM in hickory (Fig. S12C), consistent with the increased input of exudates, which have a higher C:N ratio (~40:1) than microbial necromass (~10:1 in models and 7.6:1 in global estimates), another important precursor of SOM. Thus, the increased C:N ratio of MAOM likely reflects a promotion of the direct sorption of exudate compounds to mineral surfaces (33, 34). In maple, by contrast, increasing herbivory pressure decreased the C:N ratio of MAOM, reflecting a weaker influence of herbivory on C exudation rates and on MAOM formation, and suggesting a more important role of microbial necromass for MAOM formation (i.e., greater microbial processing during MAOM formation). Although the C:N ratio of POC also differed significantly with herbivory (Fig. S12D), the slope of these relationships was comparatively shallow, suggesting a much larger influence of herbivory on MAOM.

### Implications and future directions

Our study identifies root-feeding soil fauna as an overlooked biotic control on plant C allocation to soils and reveals taxon-specific ecophysiological mechanisms through which root herbivory regulates root exudation in mature temperate forest trees. Previous work on the drivers of rhizodeposition has focused largely on direct plant responses to abiotic global change factors, yet reported exudation responses to drivers such as warming and drought vary substantially among tree species. Our results suggest that part of this variation may arise from an overlooked biotic pathway: global change drivers can alter the abundance and composition of root-feeding soil fauna, whose effects on root exudation differ among tree taxa. For example, populations of root-feeding nematodes are known to respond to environmental pressures such as warming and drought (20, 35). Consequently, variation in how root-feeder populations respond to these drivers, coupled with their taxon-specific effects on dominant tree species, could contribute to the inconsistent exudation responses previously reported under warming and drought (36–38). Belowground herbivory may therefore represent a missing mechanistic link between global change drivers and variation in tree root exudation responses.

Current frameworks for assessing the drivers of ecosystem C cycling and SOM persistence are neglecting the key component of soil fauna (36, 37). Our study moves beyond correlation-based approaches and extends field-based causal quantification of the C impact of soil animals that feed on living plant tissues, thus calling for intensifying the inclusion of soil fauna as a means to improve predictions from ecosystem C models. Results offer a basis for integrating belowground herbivory into multitrophic biogeochemistry and encourage future manipulative experiments to include additional herbivore and tree taxa and to integrate belowground herbivores with other soil fauna influencing multiple biogeochemical pathways to quantify their net effect on soil C accrual and persistence.

## Materials and Methods

### Site Descriptions and Experimental Design

In Summer 2025, we conducted a manipulative field experiment at the Griffy Woods Indiana University Research and Teaching Preserve, a temperate hardwood forest in southern Indiana, USA (86.505013°W, 39.188813°N). The study site is in a hot-summer humid continental (Köppen Dfa) climate zone, experiencing mean monthly temperatures between −1.78°C and 23.89°C and average annual precipitation of 1,254.3 mm between 1991 and 2020. The experiment used a blocked factorial design consisting of four root herbivory treatments × two dominant tree genera, with eight replicates. Manipulated root herbivory treatments included a defaunated control, ambient herbivory pressure, and ambient level increased two- and three-fold. We selected eight mature (Diameter at breast height greater than 25cm) hickory trees (seven *Carya ovata* and one *C. glabra* and eight sugar maple trees (*Acer saccharum*), representing ECM- and AM-associated trees, respectively. At each tree, four intact roots with absorptive roots (≤ 2mm diameter) connected to separate shallow woody roots were excavated, cleaned, and placed individually into a microcosm. Root herbivory treatments were established using *Pratylenchus penetrans*, a ubiquitous root-feeding nematode common across all temperate regions globally. Treatments consisted of a defaunated control (0 nematodes per microcosm), ambient (300), two-fold ambient (600), and three-fold ambient (900) herbivory levels. The ambient treatment was based on root-feeding nematode abundance measured at the study site (24). A detailed description of soil defaunation by thermal treatment (38), inoculum preparation and application methods is presented in Supplementary Methods. Microcosms were incubated underground in the field for eight weeks.

### Sample collection

Soil, root exudate, and root samples were collected after eight weeks of field incubation. Exudates were collected using an adapted cuvette collection method (39, 40) (Supplementary Methods). Exudate samples were filtered and divided for measurement of C exudation rate and metabolomic analysis. Samples for metabolomics were frozen and lyophilized, whereas samples for C exudation rate were acidified and frozen until analysis. Following exudate collection, roots were scanned using a flatbed scanner (Epson Perfection V600; Seiko Epson Corporation, Suwa, Nagano, Japan) and analyzed using RhizoVision Explorer v2.0.3 (41). We quantified root necrosis rate, mean root diameter (mm), specific root length (cm g^-1^), branching frequency (cm^-1^), and number of root tips. Root necrosis rate was calculated as the proportion of total root surface area classified as necrotic and used as an estimate of root damage. Nematodes were subsequently extracted from roots and soil to characterize post-treatment abundance (Figs. S13 and S14). Exudate collection and separation, root scanning, and nematode extraction are described in detail in Supplementary Methods.

### Root C exudation rate and metabolomics

Root C exudation rate was quantified as non-purgeable organic carbon (NPOC) using a TOC analyzer (TOC-L CSN; Shimadzu Scientific Instruments, Columbia, MD, USA) coupled to an autosampler unit (ASI-L; Shimadzu Scientific Instruments) with a volume of 15-20mL of exudate samples (Supplementary Methods). NPOC concentrations were normalized by root surface area (39, 42) and exudate collection duration. Root exudate metabolomes were characterized using liquid chromatography-mass spectrometry (LC-MS/MS) following a previously developed protocol (39) (Supplementary Methods). Analyses were conducted in positive and negative ionization modes. Metabolite concentrations were normalized to an internal standard (2-amino-3-bromo-5-methylbenzoic acid), and identified compounds were grouped into functional metabolite classes for subsequent analyses.

### SOM fractionation and CN measurement

Soils collected from the microcosms were fractionated into MAOM and POM (Supplementary Methods) for measurement of MAOC, POC, mineral-associated organic nitrogen (MAON), and particulate organic nitrogen (PON). Fractionation was performed using a particle size-based method (43). Then, subsamples of 0.15 ± 0.01 g of POM or MAOM were used to measure C and nitrogen content by high-temperature combustion using a LECO CN828 analyzer (Leco Corporation, St. Joseph, MI, USA).

### Statistical analysis

All statistical analyses were performed in R v4.3.1 (44). Prior to analysis, two outliers in root C exudation rate and two in root necrosis rate were identified using Rosner’s test and removed using the *EnvStats* package (45). Linear mixed-effects models implemented with the *nlme* package (46, 47) were used to test the effects of root herbivory level, tree type, and their interaction on root C exudation rate, root necrosis rate, MAOC, POC, and MAOM and POM C ratios. Tree DBH was included as a random effect. Marginal and conditional R^2^, representing variance explained by fixed effects and by fixed and random effects combined, respectively, were calculated using the *piecewiseSEM* package (48, 49).

Metabolite concentrations were normalized to an internal standard and autoscaled using the *mdatools* package (50). Compound richness and concentrations of metabolite functional groups were analyzed using linear mixed-effects models as described above. Differences in overall exudate composition among herbivory levels and tree types were assessed by PERMANOVA with 999 permutations and visualized using non-metric multidimensional scaling (NMDS). Both analyses were based on Bray–Curtis dissimilarities and conducted using the *vegan* package (51, 52). Individual metabolites associated with herbivory treatments were identified by indicator species analysis using the *indicspecies* package (53), with significance assessed by 999 permutations. The indicator value (IndVal) combines specificity, the degree to which a metabolite is uniquely associated with a treatment group, and sensitivity, the frequency with which it occurs within that group. IndVal ranges from 0 to 1, with values approaching 1 indicating a strong association with a treatment. Statistical significance was assessed using 999 permutations, and metabolites with p < 0.05 were considered indicators of their respective herbivory treatment.

Structural equation models (SEMs) implemented in *piecewiseSEM* (48) were used to assess direct and indirect pathways linking root herbivory to MAOC through root necrosis, C exudation rate, and exudate composition. We focused on MAOC because its formation is closely linked to mineral association of low-molecular-weight root-derived compounds, whereas POC formation is primarily associated with particulate plant inputs (54, 55). Separate SEMs were constructed for hickory and maple to evaluate tree-specific pathways. Root exudate composition was represented by a composite variable derived from metabolite functional groups selected through exploratory factor analysis using the *psych* package (56). The suitability of functional groups for inclusion was evaluated using the Kaiser–Meyer–Olkin measure of sampling adequacy (57, 58) and Bartlett’s test of sphericity (59) with the *psych* package. Functional groups meeting these criteria were subjected to exploratory factor analysis (60, 61), from which 6–7 groups were retained to represent root exudate composition. Standardized path coefficients (β) and associated p-values were used to evaluate the strength and statistical support of relationships among variables.

## Acknowledgments

We thank Navya Ayitha, Gwendolyn Lloyd, Felipe Barros, and Young-Eui Oh for their assistance with the field experiment, Matthew Taylor for soil CN measurement, and Jacob Scharfetter for exudate metabolomics analysis. This work was partially funded through a Fischer Faculty Fellowship to ALCF.

## References

1. D. B. Metcalfe, et al., Herbivory makes major contributions to ecosystem carbon and nutrient cycling in tropical forests. Ecology Letters 17, 324–332 (2014).

2. J. Å. Kristensen, J. Rousk, D. B. Metcalfe, Below-ground responses to insect herbivory in ecosystems with woody plant canopies: A meta-analysis. Journal of Ecology 108, 917–930 (2020).

3. H. Gan, K. Wickings, Root herbivory and soil carbon cycling: Shedding “green” light onto a “brown” world. Soil Biology and Biochemistry 150, 107972 (2020).

4. R. B. Jackson, et al., The Ecology of Soil Carbon: Pools, Vulnerabilities, and Biotic and Abiotic Controls. Annual Review of Ecology, Evolution, and Systematics 48, 419–445 (2017).

5. S. H. Villarino, P. Pinto, R. B. Jackson, G. Piñeiro, Plant rhizodeposition: A key factor for soil organic matter formation in stable fractions. Science Advances 7, eabd3176 (2021).

6. A. L. C. Franco, L. A. Gherardi, “Chapter Three - Soil fauna shape plant productivity and its belowground allocation across global grasslands” in Advances in Agronomy, D. L. Sparks, Ed. (Academic Press, 2025), pp. 109–132.

7. G. W. Yeates, S. Saggar, C. S. Denton, C. F. Mercer, Impact of Clover Cyst Nematode (Heterodera Trifolii) Infection On Soil Microbial Activity in the Rhizosphere of White Clover (Trifolium Repens) - a Pulse-Labelling Experiment. (1998). 10.1163/005225998X00082.

8. C. S. Denton, R. D. Bardgett, R. Cook, P. J. Hobbs, Low amounts of root herbivory positively influence the rhizosphere microbial community in a temperate grassland soil. Soil Biology and Biochemistry 31, 155–165 (1998).

9. Bardgett, Denton, Cook, Below-ground herbivory promotes soil nutrient transfer and root growth in grassland. Ecology Letters 2, 357–360 (1999).

10. N. R. Chari, et al., Estimating the global root exudate carbon flux. Biogeochemistry 167, 895–908 (2024).

11. C. Fossum, et al., Belowground allocation and dynamics of recently fixed plant carbon in a California annual grassland. Soil Biology and Biochemistry 165, 108519 (2022).

12. N. W. Sokol, Sara. E. Kuebbing, E. Karlsen-Ayala, M. A. Bradford, Evidence for the primacy of living root inputs, not root or shoot litter, in forming soil organic carbon. New Phytologist 221, 233–246 (2019).

13. N. W. Sokol, J. Sanderman, M. A. Bradford, Pathways of mineral-associated soil organic matter formation: Integrating the role of plant carbon source, chemistry, and point of entry. Global Change Biology 25, 12–24 (2019).

14. N. R. Chari, B. N. Taylor, Soil organic matter formation and loss are mediated by root exudates in a temperate forest. Nat. Geosci. 15, 1011–1016 (2022).

15. I. A. Shabtai, et al., Root exudates simultaneously form and disrupt soil organo-mineral associations. Commun Earth Environ 5, 699 (2024).

16. T. Wen, et al., Root exudate chemistry affects soil carbon mobilization via microbial community reassembly. Fundamental Research 2, 697–707 (2022).

17. L. Chen, Y. Liu, The Function of Root Exudates in the Root Colonization by Beneficial Soil Rhizobacteria. Biology (Basel) 13, 95 (2024).

18. M. J. Zwetsloot, A. Kessler, T. L. Bauerle, Phenolic root exudate and tissue compounds vary widely among temperate forest tree species and have contrasting effects on soil microbial respiration. New Phytologist 218, 530–541 (2018).

19. S. Haase, L. Ruess, G. Neumann, S. Marhan, E. Kandeler, Low-level herbivory by root-knot nematodes (Meloidogyne incognita) modifies root hair morphology and rhizodeposition in host plants (Hordeum vulgare). Plant Soil 301, 151–164 (2007).

20. A. L. C. Franco, et al., Drought suppresses soil predators and promotes root herbivores in mesic, but not in xeric grasslands. Proceedings of the National Academy of Sciences 116, 12883–12888 (2019).

21. E. A. Shaw, C. M. Boot, J. C. Moore, D. H. Wall, J. S. Baron, Long-term nitrogen addition shifts the soil nematode community to bacterivore-dominated and reduces its ecological maturity in a subalpine forest. Soil Biology and Biochemistry 130, 177–184 (2019).

22. J. Zhao, et al., Sentinel soil invertebrate taxa as bioindicators for forest management practices. Ecological Indicators 24, 236–239 (2013).

23. A. Afzal, T. Mukhtar, Revolutionizing nematode management to achieve global food security goals - An overview. Heliyon 10, e25325 (2024).

24. K. C. Ball, et al., Seasonal variation of micro and macro soil invertebrate communities in a temperate forest. Applied Soil Ecology 218, 106729 (2026).

25. A. Canarini, C. Kaiser, A. Merchant, A. Richter, W. Wanek, Root Exudation of Primary Metabolites: Mechanisms and Their Roles in Plant Responses to Environmental Stimuli. Front. Plant Sci. 10 (2019).

26. C. Ristok, et al., Plant species richness elicits changes in the metabolome of grassland species via soil biotic legacy. Journal of Ecology 107, 2240–2254 (2019).

27. M. Oss, A. Kruve, K. Herodes, I. Leito, Electrospray Ionization Efficiency Scale of Organic Compounds. Anal. Chem. 82, 2865–2872 (2010).

28. A. Kruve, K. Kaupmees, J. Liigand, I. Leito, Negative Electrospray Ionization via Deprotonation: Predicting the Ionization Efficiency. Anal. Chem. 86, 4822–4830 (2014).

29. J. Bergmann, et al., The fungal collaboration gradient dominates the root economics space in plants. Science Advances 6, eaba3756 (2020).

30. A. Weigelt, et al., An integrated framework of plant form and function: the belowground perspective. New Phytologist 232, 42–59 (2021).

31. A. Williams, F. T. de Vries, Plant root exudation under drought: implications for ecosystem functioning. New Phytologist 225, 1899–1905 (2020).

32. F. el Z. Haichar, C. Santaella, T. Heulin, W. Achouak, Root exudates mediated interactions belowground. Soil Biology and Biochemistry 77, 69–80 (2014).

33. S. Manzoni, A. Porporato, Soil carbon and nitrogen mineralization: Theory and models across scales. Soil Biology and Biochemistry 41, 1355–1379 (2009).

34. M. Rötzer, et al., Coupled C and N turnover in a dynamic pore scale model reveal the impact of exudate quality on microbial necromass formation. SOIL 12, 205–226 (2026).

35. K. E. Mueller, et al., Elevated CO2 and warming shift the functional composition of soil nematode communities in a semiarid grassland. Soil Biology and Biochemistry 103, 46–51 (2016).

36. J. D. M. Lejoly, K. Mason-Jones, G. F. C. Veen, A soil food web approach to integrate soil fauna into multitrophic biogeochemistry. Commun Earth Environ 7, 322 (2026).

37. A. S. Grandy, W. R. Wieder, K. Wickings, E. Kyker-Snowman, Beyond microbes: Are fauna the next frontier in soil biogeochemical models? Soil Biology and Biochemistry 102, 40–44 (2016).

38. A. L. C. Franco, et al., Nematode exclusion and recovery in experimental soil microcosms. Soil Biology and Biochemistry 108, 78–83 (2017).

39. B. D. Hafner, O. Pietz, W. L. King, J. B. Scharfetter, T. L. Bauerle, Early developmental shifts in root exudation profiles of five Zea mays L. genotypes. Plant Science 354, 112439 (2025).

40. R. P. Phillips, Y. Erlitz, R. Bier, E. S. Bernhardt, New approach for capturing soluble root exudates in forest soils. Functional Ecology 22, 990–999 (2008).

41. A. Seethepalli, et al., RhizoVision Explorer: open-source software for root image analysis and measurement standardization. AoB PLANTS 13, plab056 (2021).

42. M. Brunn, et al., Carbon allocation to root exudates is maintained in mature temperate tree species under drought. New Phytologist 235, 965–977 (2022).

43. C. A. Cambardella, E. T. Elliott, Particulate Soil Organic-Matter Changes across a Grassland Cultivation Sequence. Soil Science Society of America Journal 56, 777–783 (1992).

44. R Core Team, R: A Language and Environment for Statistical Computing. (2023).

45. S. P. Millard, EnvStats: An R Package for Environmental Statistics (Springer, 2013).

46. J. C. Pinheiro, D. M. Bates, R Core Team, nlme: Linear and Nonlinear Mixed Effects Models. (2024). Deposited 14 August 2024.

47. J. C. Pinheiro, D. M. Bates, Mixed-Effects Models in S and S-PLUS (Springer-Verlag, 2000).

48. J. S. Lefcheck, piecewiseSEM: Piecewise structural equation modelling in r for ecology, evolution, and systematics. Methods in Ecology and Evolution 7, 573–579 (2016).

49. S. Nakagawa, H. Schielzeth, A general and simple method for obtaining R2 from generalized linear mixed-effects models. Methods in Ecology and Evolution 4, 133–142 (2013).

50. S. Kucheryavskiy, mdatools – R package for chemometrics. Chemometrics and Intelligent Laboratory Systems 198, 103937 (2020).

51. M. J. Anderson, A new method for non-parametric multivariate analysis of variance. Austral Ecology 26, 32–46 (2001).

52. J. Oksanen, et al., vegan: Community Ecology Package. (2026). Deposited 4 March 2026.

53. M. D. Cáceres, P. Legendre, Associations between species and groups of sites: indices and statistical inference. Ecology 90, 3566–3574 (2009).

54. J. Lehmann, M. Kleber, The contentious nature of soil organic matter. Nature 528, 60–68 (2015).

55. J. M. Lavallee, J. L. Soong, M. F. Cotrufo, Conceptualizing soil organic matter into particulate and mineral-associated forms to address global change in the 21st century. Global Change Biology 26, 261–273 (2020).

56. W. Revelle, psych: Procedures for Psychological, Psychometric, and Personality Research. (2026). Deposited 16 May 2026.

57. H. F. Kaiser, A Second Generation Little Jiffy. Psychometrika 35, 401–415 (1970).

58. H. F. Kaiser, J. Rice, Little Jiffy, Mark Iv. Educational and Psychological Measurement 34, 111–117 (1974).

59. M. S. Bartlett, Tests of Significance in Factor Analysis. British Journal of Statistical Psychology 3, 77–85 (1950).

60. L. R. Fabrigar, D. T. Wegener, R. C. MacCallum, E. J. Strahan, Evaluating the use of exploratory factor analysis in psychological research. Psychological Methods 4, 272–299 (1999).

61. D. Lüdecke, et al., parameters: Processing of Model Parameters. (2026). Deposited 9 May 2026.

