## Supplementary information for "Belowground Herbivory Shapes Carbon Release from Tree Roots and Its Retention in Soil"

* Jaeyoung Yoo, Andre L.C. Franco

**This PDF file includes:**

Supplementary Methods

Figures S1 to S14

Tables S1 to S2

SI References

**Supplementary Methods**

**Soil defaunation, inoculum preparation and application.** We used a 10cm diameter hole cutter to collect ~50 kg of topsoil (10 cm depth) from the root zone of the sixteen selected trees and homogenized it separately within hickory plots and maple plots. To remove native soil fauna, soils were thermally treated following (1) For defaunation, air-dried, sieved soil (6.25 mm) was placed into aluminum dishes with small holes at the bottom, ensuring a depth of 5 cm. Soil was then pre-wetted by gently spraying the surface until water passed through the holes in the bottom of the trays. The pre-wetted soil was left at 4 °C for 24 h. Next, the trays were transferred to an oven set at 65 °C for 48 h. This thermal treatment eliminates 99% of the nematode population and suppresses recolonization for at least 8 weeks (1). We subsampled 10 aliquots of defaunated soil and found no living nematodes after running Baermann funnel extraction for 72 hours (2, 3). Our experimental units consisted of customized microcosms made with 5×9" Whirl-Pak bags in which we opened a 3×5" square on one side and covered the opening with a 25-micron mesh. This was to allow movement of air and water while blocking nematodes and other invertebrate fauna from entering or exiting, thus reducing microclimate variation within the microcosm while maintaining experimental treatments as intended. Each microcosm was filled with 700 g of rewetted defaunated soil. Soil moisture was adjusted to an average ambient level of 25.88 ± 0.02% reported by (4) at the same site. Accordingly, each microcosm received 520 g of defaunated soil plus 180 g of deionized water, resulting in a soil moisture content of 25.71%, closely matching ambient soil moisture at the site.

In June 2025, our team carefully excavated the topsoil around each selected tree to identify intact roots. At each tree, we identified four roots, including absorptive roots located approximately 50cm from the trunk (i.e., 2 mm or less in diameter), each branching from separate shallow woody roots which could be clearly tracked back to the selected tree. Once identified, roots were carefully but extensively rinsed with deionized water, and the top part (~6 to 8 inches) placed into the microcosms containing 700 g of rewetted defaunated soil prior to inoculation with nematode treatments. To establish a realistic baseline for root herbivory treatments, we used a mean ambient herbivory pressure of 34.75 ± 8.06 endoparasitic root-feeding nematodes per 100 g fresh soil obtained in a previous study in Spring 2023 at the same site (4). Accordingly, microcosms receiving the ambient herbivory treatment were inoculated with 300 root-feeding nematodes, or approximately 40 nematodes per 100g fresh soil. We used Pratylenchus penetrans, a migratory endoparasitic root-feeding nematode, from laboratory cultures to establish a gradient from low to high root herbivory pressure. This nematode species is found in all temperate regions globally (5). P. penetrans cultures grew in Petri dishes with corn roots and Gamborg B5 (GB5) medium, as described in (6), and inoculum was prepared and applied as described in (7). Briefly, we extracted *P. penetrans* from 8 Petri dishes by slicing the culture medium and corn roots and placing them on Baermann funnels for 24 hr. The extracted solutions were combined in a 1,000 mL flask, and nematodes were counted in ten 1-mL aliquots to obtain an average nematode density in the inoculum, which was 79.10 ± 3.82 individuals mL^-1^, consisting of all vermiform stages in water. Thus, the volume of *P. penetrans* inoculum added to achieve the ambient herbivory treatment (hereafter called “low herbivory treatment”) was 3.8 mL. We then produced a gradient of increased root herbivory pressure by applying treatments with two and three times the number of root herbivores inoculated in the low herbivory treatment. The mid-level and high herbivory treatments received 7.6 mL (or 600 nematodes) and 11.4 mL (or 900 nematodes) of P. penetrans inoculum, respectively. To keep the total inoculum volume constant across treatments and ensure equivalent water and nutrient inputs, we prepared a nematode-free nutrient solution by adding the contents of Petri dishes containing GB5 medium and corn roots but no nematodes to additional Baermann funnels. The nutrient inoculum was extracted following the same procedure as the P. penetrans inoculum. No nematodes were detected in the nutrient inoculum. The low, mid, and high herbivory treatments received 11.2, 7.4, and 3.6 mL of nutrient inoculum, respectively, thereby standardizing the applied volume of inoculum at 15 mL across all treatments. A defaunated control treatment received 15 mL of nematode-free nutrient inoculum (i.e., 0 P. penetrans per microcosm). All inocula were stored at 4°C for 24–48 hours, brought to the field site inside a cooler containing ice packs, and inoculated with 5-ml glass measuring pipettes attached to a pipet controller (Fisherbrand^®^). Those solutions were constantly and gently homogenized during inoculation. At each microcosm, the solutions were slowly poured into the soil inside the bags. A total of 64 microcosms were incubated underground in the field for eight weeks and protected with plastic mesh and landscape staples throughout the incubation period.

**Adapted cuvette collection method.** First, the microcosms containing intact tree roots were carefully opened, and the roots were removed from the soil and gently rinsed with ultrapure water to remove adhering soil particles. The top part of intact roots from each microcosm was placed into a 30 mL plastic syringe (Luer Lock Syringes; Air-Tite Products, Virginia Beach, VA, USA), and glass beads were added (750 μm diameter; La De Da Designs, Baton Rouge, LA, USA). Roots were then flushed three times before being filled with ultrapure water and sealed at the top of each syringe with Parafilm. The water-filled syringes were left in the field for 24 hours wrapped in aluminum foil. A syringe containing glass beads but no roots was placed next to each root exudate collection syringe to serve as a blank control and was also left for 24 hours wrapped in aluminum foil to account for potential contamination introduced during syringe setup. After 24 hours, roots were finally detached from trees and cut near the top of the syringe before being transported to the laboratory. Soil samples and exudate syringes were stored at 4°C until processing or analysis.

**Exudate collection and separation, root scanning, and nematode extraction.** On the same day the syringes with root exudate were collected, 20 mL of ultrapure water was added to each syringe in the laboratory to ensure complete recovery of the solution remaining in the syringe, while extracted into acid-washed glass vials (VOA Sampling Vials; RESTEK, Bellefonte, PA, USA) using a vacuum pump. Exudates were filtered through sterile polyvinylidene fluoride (PVDF) syringe filters (Millex PVDF, pore size 0.22 μm, diameter 33 mm, sterile, hydrophilic; Millex, Duluth, GA, USA) and divided into two vials: 10 mL for metabolomics analysis and the remainder for root C exudation rate measurement. Exudate vials designated for metabolomics were immediately frozen at −20°C and lyophilized. Vials designated for root C exudation rate measurements were acidified with 1–2 drops of 12 M HCl and frozen at −20°C until further analysis. All consumables used to collect and store root exudates had been previously acid-washed in 10% nitric acid or cleaned with C-free detergent (Liquinox; ALCONOX, White Plains, NY, USA).

Following exudate collection, roots remaining in the syringes were rinsed with reverse osmosis (RO) water, scanned using a flatbed scanner (Epson Perfection V600; Seiko Epson Corporation, Suwa, Nagano, Japan) at 1,200 dpi in grayscale, and analyzed for root necrosis rate, mean root diameter (mm), specific root length (cm g^-1^), branching frequency (cm^-1^), and number of root tips using RhizoVision Explorer Version 2.0.3 (8). Root necrosis rate was calculated as the necrotic root surface area divided by the total root surface area and was used as an estimate of root damage caused by nematodes. Necrotic tissue appears significantly brighter in scanned images, enabling detection by applying different image thresholding levels in RhizoVision Explorer. Image thresholding levels of 80 and 130 were used to capture root necrosis surface area and total root surface area, respectively. Following scanning, roots were cut into 1- 2 cm pieces, and nematodes were extracted from the cut roots using Baermann funnels and 72h extraction time (2, 9) and were fixed with 5% formalin for further identification (10). Then, all roots were dried at 65°C for 48 hours to determine their dry mass. Soil nematodes were extracted from 100g of fresh soil and fixed as roots for further identification. Post-treatment abundance of nematodes in roots and soils is presented in Fig. S13 and S14.

**Root C exudation rate measurement.** A volume of 15–20 mL of each exudate sample was analyzed for non-purgeable organic carbon (NPOC) using a TOC analyzer (TOC-L CSN; Shimadzu Scientific Instruments, Columbia, MD, USA) coupled to an autosampler unit (ASI-L; Shimadzu Scientific Instruments). Frozen samples were thawed at room temperature for 24 hours prior to analysis. A six-point calibration curve (1, 2, 5, 10, 25, and 50 mg NPOC L⁻¹) was established before analysis. For quality control, an external certified reference standard (5 and 10 mg NPOC L⁻¹; Ricca Chemical Company, Arlington, TX, USA) and an internal laboratory standard (5 and 10 mg NPOC L⁻¹) were used to validate measurement precision. Measured NPOC concentrations were normalized to root surface area and the duration of root exudate collection.

**LC-MS analysis and data processing.** Lyophilized exudates were redissolved with 10 mL of 4°C methanol (Millipore-Sigma, Burlington, MA, USA) and each sample was subsequently vortexed for 2 minutes. Samples were spiked with 0.100 mM of 2-amino-3-bromo-5-methylbenzoic acid (internal standard) and vortexed again for 1 minute. The exudates were analyzed using LC-MS/MS following a previously developed protocol (11). Chromatography was performed on a Dionex UltiMate 3000 UPLC stack (Thermo-Fisher, Waltham, MA, USA) equipped with an Ultimate 3000 RS pump, Ultimate 3000 RS autosampler, Ultimate 3000 RS Column Compartment, and Ultimate 3000 RS diode array detector (DAD). The LC-Stack was coupled to a Thermo Q-Exactive Hybrid Quadrupole Orbitrap mass spectrometer. Chromatographic separation was achieved using TitanTM C18 UHPLC reverse phase column [2.1 mm x 100 mm (1.9μm), Agilent Technologies] with a flow rate of 0.5 mL min^-1^ and a column temperature of 40°C. The mobile phases consisted of a 0.1% formic acid solution mixed with Optima (Thermo-Fisher) LC-MS grade water (eluent A) and a 0.1% formic acid solution mixed with Optima LC-MS grade acetonitrile (eluent B). The gradient consisted of 0 to 15 min, 98% to 5% A; 15 to 15.5 min hold at 5% A; 15.5 to 15.6 min, 5% to 98% A; and 15.6 to 16.0 min hold at 98% A. The injection volume was 5 μL. Mass spectral parameters for both positive and negative modes were set at a spray voltage of 3500 V; capillary temperature of 380°C; sheath gas of 60 (arbitrary units); auxiliary gas of 20 (arbitrary units); and a HESI probe temperature of 300°C. Full-scan mass spectra were detected under a resolution 70,000 full width at half maximum and with a mass range m/z 70-1050 for both negative and positive ionization modes. Spectral acquisition was between 0.3–15 min. For MS2 acquisition, collision energies of 10, 20, and 40 V were used. Fragments comprising MS2 contributions were grouped based on having mass tolerance under 2.5 ppm, 0.05 RT tolerance, and a minimum of three area contributions.

Mass spectral data were pre-processed (baseline correction and checked for contamination) using Xcalibur 4.3 (Thermo-Fisher, Waltham, MA, USA). Identification and quantification across samples were conducted in Compound Discoverer 3.4 SP1 (Thermo-Fisher) using an untargeted metabolomic workflow. For peak detection, a mass tolerance of 5 ppm, S/N threshold of 3, and a minimum peak intensity of 10000 (arbitrary units) were used. Quality scores were applied using mzLogic, and annotations were garnered from these spectral libraries: Bennett Group at SUNY Oneonta (BLONDES); Carotenoids Database; ChemBank; Chemical Biology Department, Max Planck Institute of Molecular Physiology; E. coli Metabolome Database; FDA; Food and Agriculture Organization of the United Nations; Indofine; KEGG; LipidMAPS; MassBank; Nature Chemical Biology; Phenol-Explorer; Sequoia Research Products; Shaw lab at Southern Illinois University Edwardsville; Yeast Metabolome Database. Additionally, m/z values and subsequent annotations were compared to an in-house tree exudate library consisting of 540 standards. Lastly, quality control batch corrections were then applied.

**Soil fractionation.** Briefly, 5.25–5.75 g of 2 mm-sieved, air-dried soil was additionally dried overnight at 60°C. The soil was then dispersed in 25 mL of 0.5% sodium hexametaphosphate solution by shaking on a reciprocal shaker at 112 rpm for 18 hours. After shaking, the dispersed soil suspension was manually passed through a 53 μm sieve and rinsed with RO water until the water ran clear. The soil material retained on top of the sieve and that which passed through were each transferred to separate aluminum soil canisters, dried at 60°C for one week, and ground to a fine powder using a roller mill with 2–3 rods per sample to minimize variability in measured C due to differences in particle size (12, 13). Material retained on top of the sieve was analyzed for POM content, while material passing through the sieve was analyzed for MAOM content.

Figures

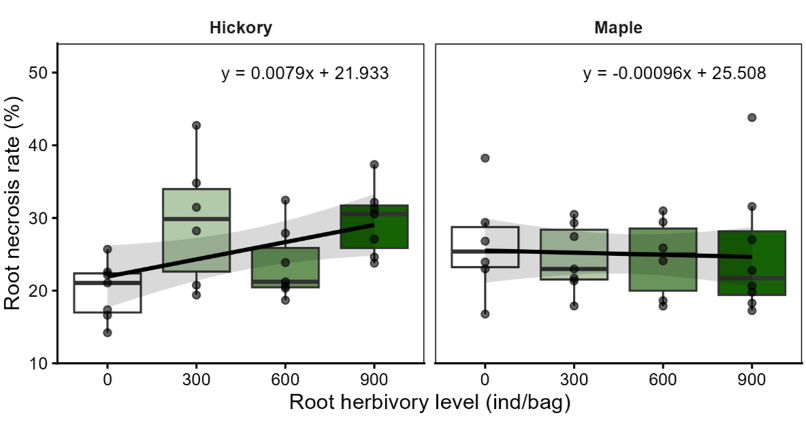

Fig. S1. Root necrosis rates (%) across root herbivory levels (nematodes density per microcosm) in hickory and maple plots (n=54). Boxes represent the interquartile range (IQR), with the center line indicating the median. Whiskers extend to 1.5 × IQR, and individual points represent biological replicates. Darker green shading indicates higher herbivory levels. The solid line represents a linear regression fit with 95% confidence intervals (shaded area).

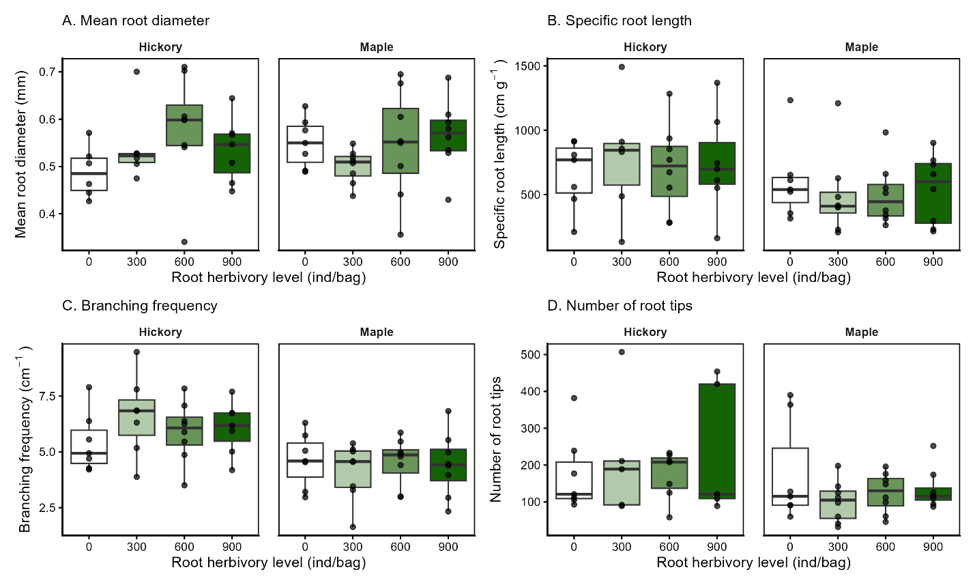

Fig. S2. Root morphological traits across root herbivory levels (0, 300, 600, and 900 nematodes per microcosm) in hickory and maple plots: (A) mean root diameter (mm), (B) specific root length (cm g^-1^), (C) branching frequency (cm^-1^), and (D) number of root tips. Boxes represent the IQR, with the centerline indicating the median. Whiskers extend to 1.5 × IQR, and individual points denote raw observations. Color intensity reflects herbivory level, from light (low) to dark (high).

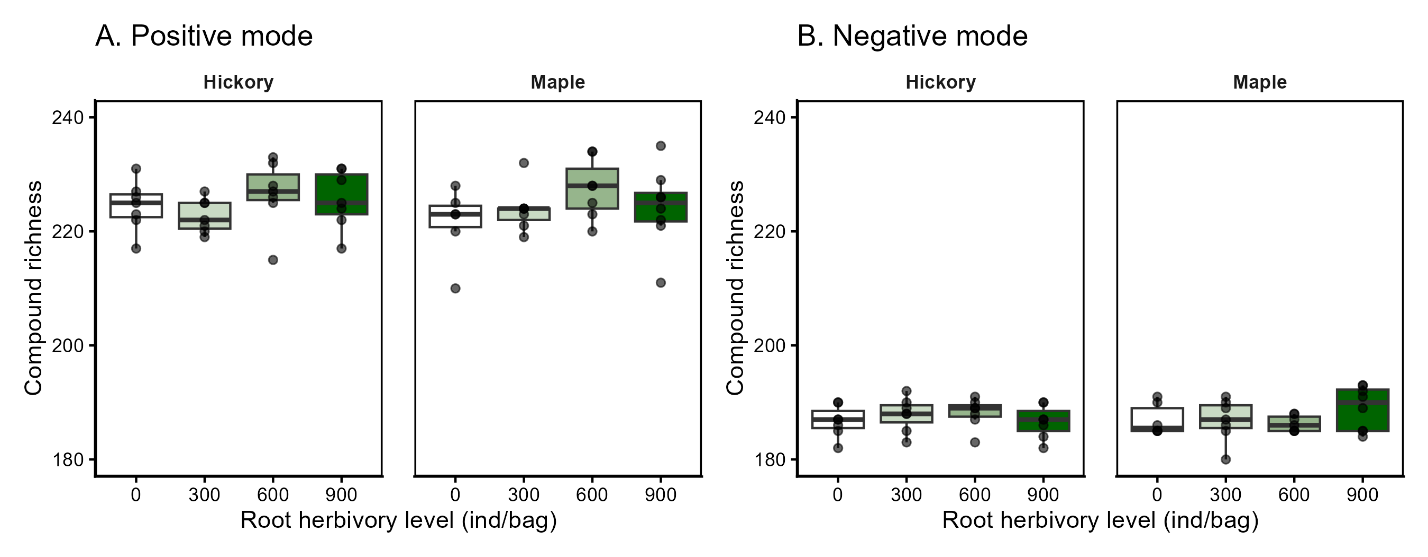
**Fig. S3.** Compound richness of root exudates across root herbivory levels (0, 300, 600, and 900 nematodes per microcosm) in hickory and maple plots in (A) positive and (B) negative ionization modes (n=56). Boxes represent the interquartile range (IQR), with the centerline indicating the median. Whiskers extend to 1.5 × IQR, and individual points denote raw observations. Color intensity reflects herbivory level, from light (low) to dark (high).

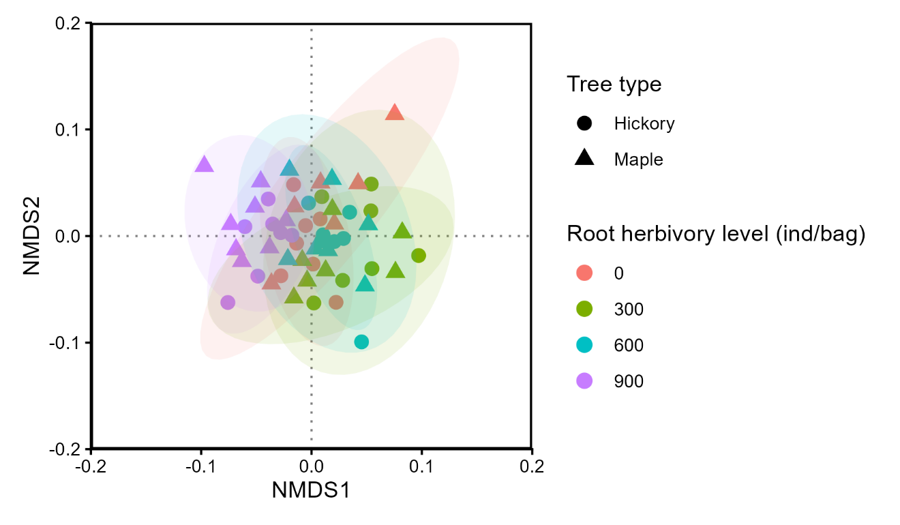

Fig. S4. Non-metric multidimensional scaling (NMDS) ordination of root exudate composition in hickory and maple plots across root herbivory levels (0, 300, 600, and 900 nematodes per microcosm) in positive ionization modes. Each point represents one biological replicate (n = 56). Symbols denote tree type (circle = hickory; triangle = maple) and colors indicate herbivory level (red = 0, green = 300, teal = 600, purple = 900 nematodes per bag). Ellipses represent 95% confidence intervals around group centroids by herbivory level.

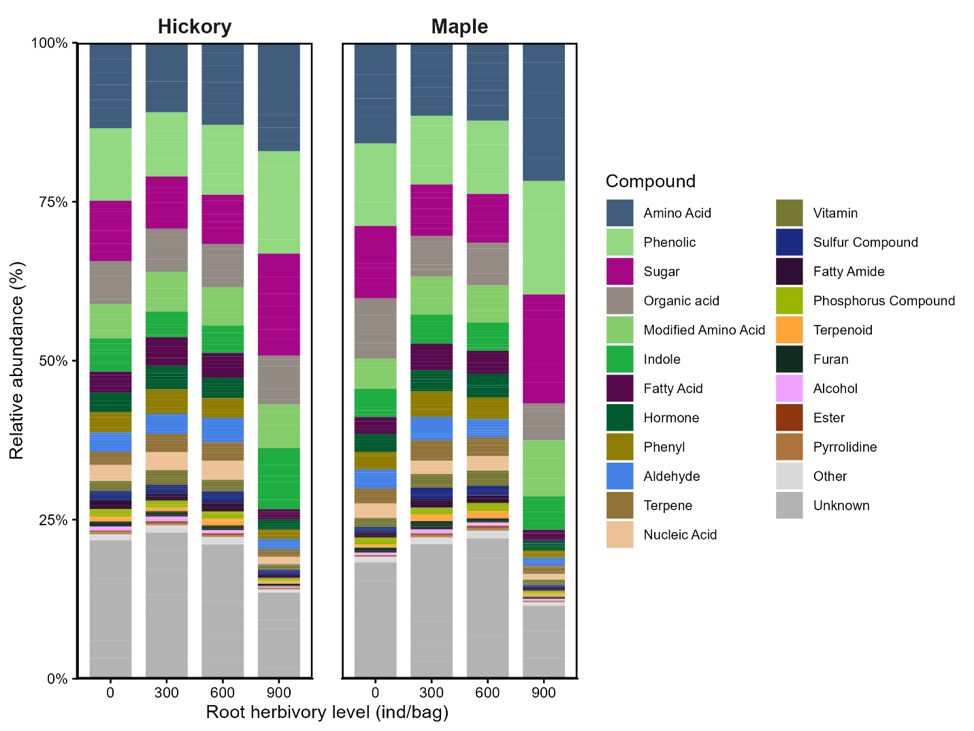

Fig. S5. Relative abundance (%) of root exudate compound classes across root herbivory levels (0, 300, 600, and 900 nematodes per microcosm) in hickory and maple plots in positive ionization modes (n = 56). Each bar represents the mean relative abundance of identified compound classes per herbivory level. Colors denote compound classes as indicated in the legend. Gray bars represent compounds of unknown classification.

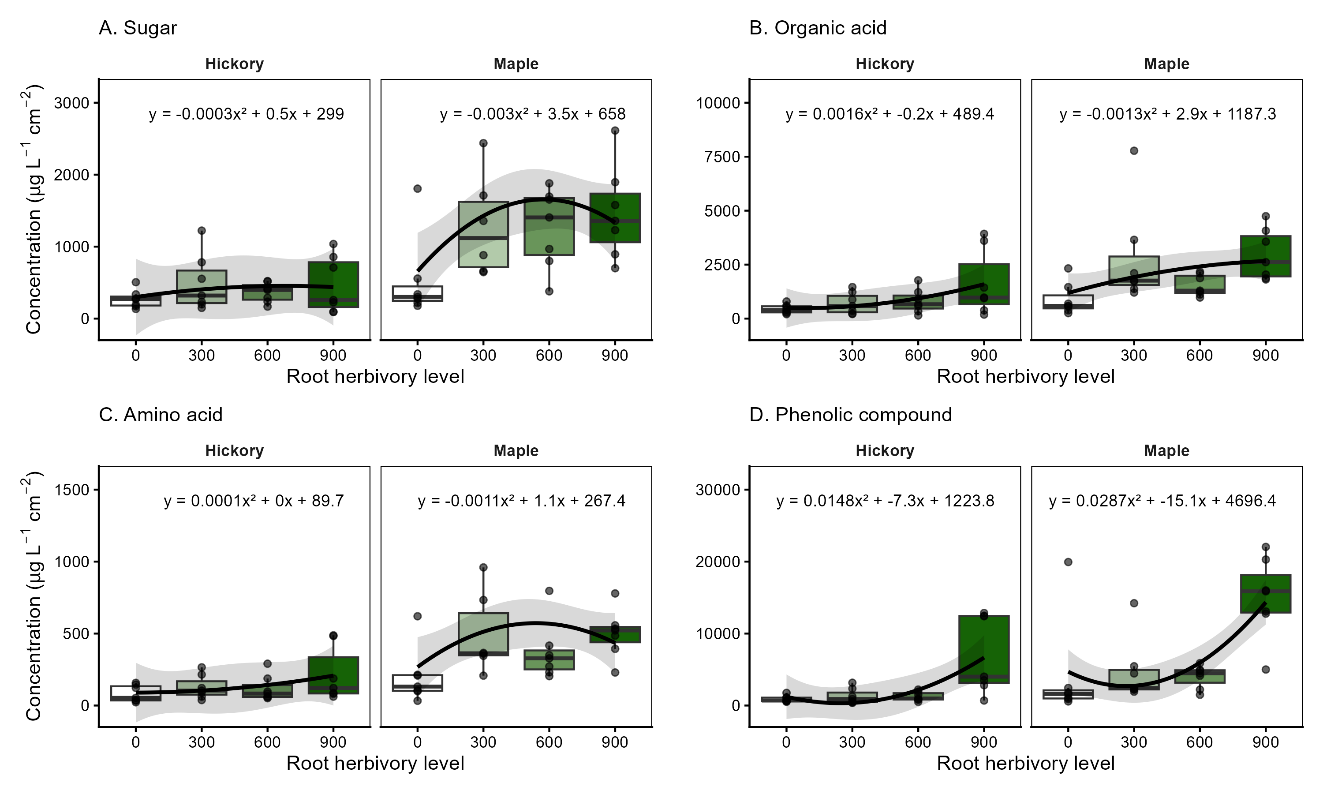

Fig. S6. Concentrations standardized by root area (μg L^-1^ cm^-2^) of (A) sugars, (B) organic acids, (C) amino acids, and (D) phenolic compounds in root exudates across root herbivory levels (0, 300, 600, and 900 nematodes per microcosm) in hickory and maple plots (negative ionization mode) (n=56). Boxes represent the interquartile range (IQR), with the center line indicating the median. Whiskers extend to 1.5 × IQR, and individual points represent biological replicates. Darker green shading indicates higher herbivory levels. The solid line represents a linear regression fit with 95% confidence intervals (shaded area).

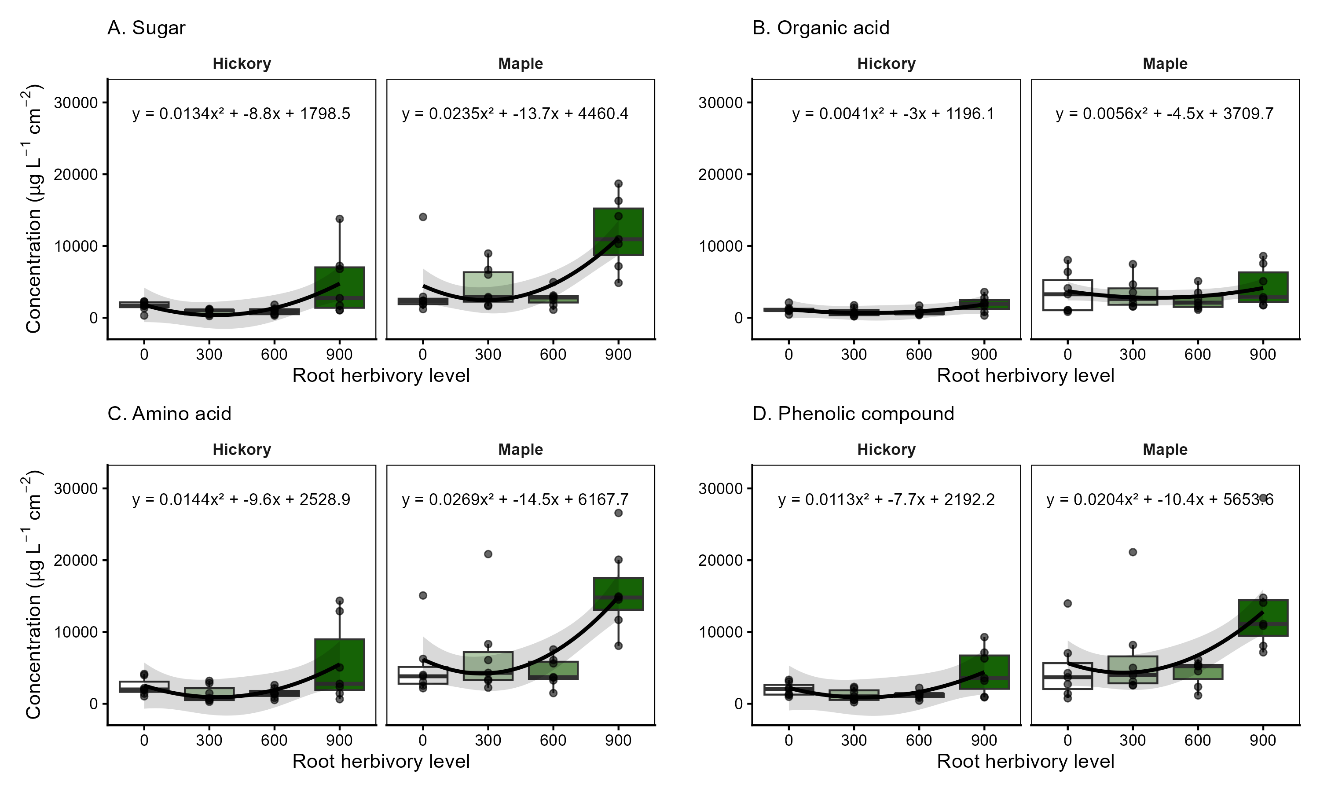

Fig. S7. Concentrations standardized by root area (μg L^-1^ cm^-2^) of (A) sugars, (B) organic acids, (C) amino acids, and (D) phenolic compounds in root exudates across root herbivory levels (0, 300, 600, and 900 nematodes per microcosm) in hickory and maple plots (positive ionization mode) (n=56). Boxes represent the interquartile range (IQR), with the center line indicating the median. Whiskers extend to 1.5 × IQR, and individual points represent biological replicates. Darker green shading indicates higher herbivory levels. The solid line represents a linear regression fit with 95% confidence intervals (shaded area).

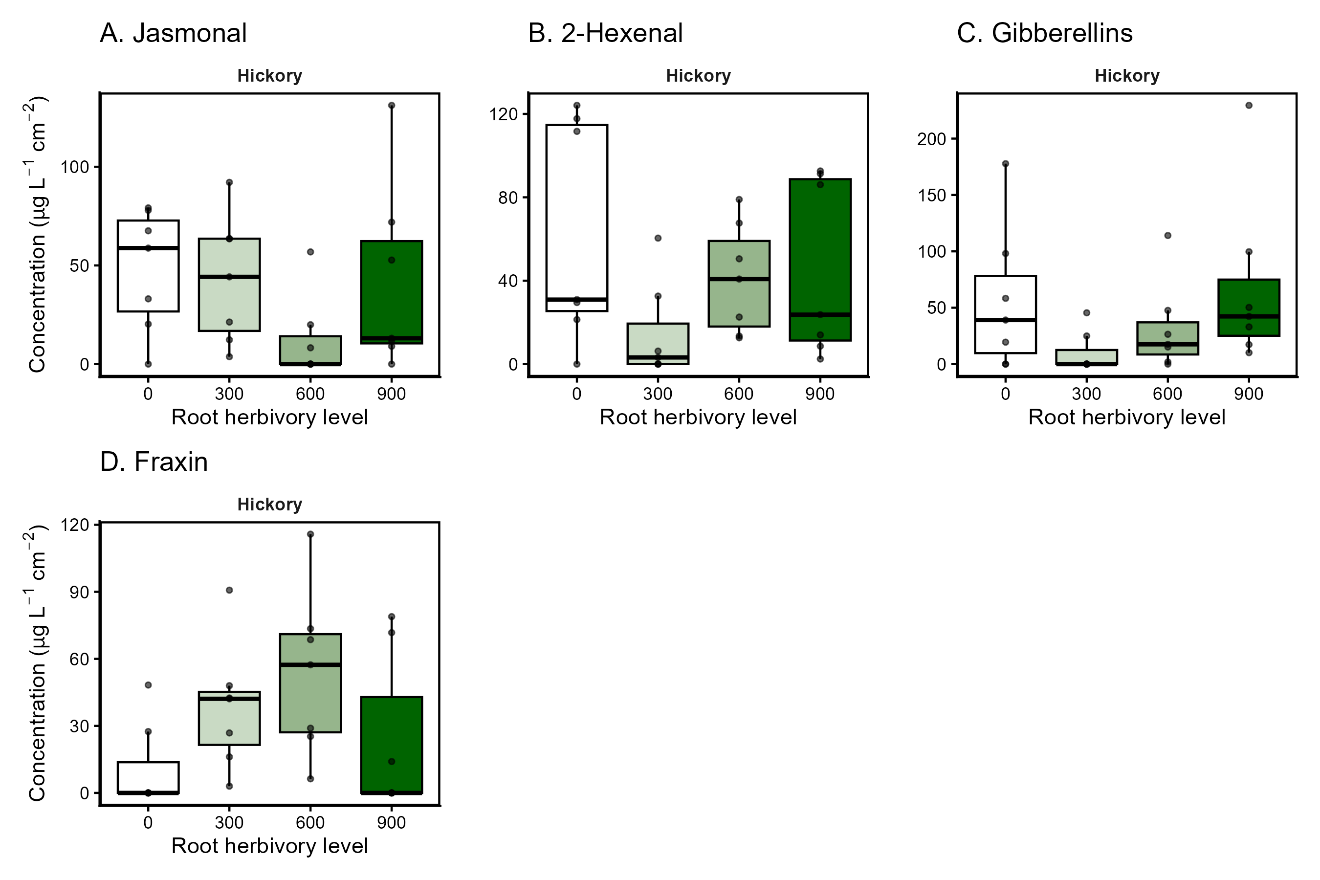

Fig. S8. Concentrations standardized by root area (μg L^-1^ cm^-2^) of individual indicator compounds identified in hickory plots across root herbivory levels (0, 300, 600, and 900 nematodes per microcosm) in positive ionization mode: (A) Jasmonal, (B) 2-Hexenal, (C) Gibberellins, and (D) Fraxin (n=56). Boxes represent the IQR, with the centerline indicating the median. Whiskers extend to 1.5 × IQR, and individual points denote raw observations. Color intensity reflects herbivory level, from light (low) to dark (high).

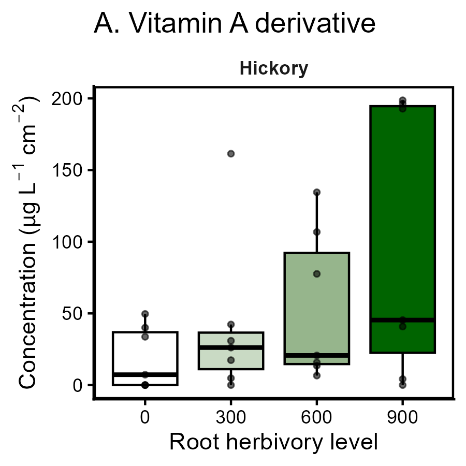

Fig. S9. Concentrations standardized by root area (μg L^-1^ cm^-2^) of the individual indicator compound identified in hickory plots across root herbivory levels (0, 300, 600, and 900 nematodes per microcosm) in negative ionization mode: (A) Vitamin A derivative (n=56). Boxes represent the IQR, with the centerline indicating the median. Whiskers extend to 1.5 × IQR, and individual points denote raw observations. Color intensity reflects herbivory level, from light (low) to dark (high).

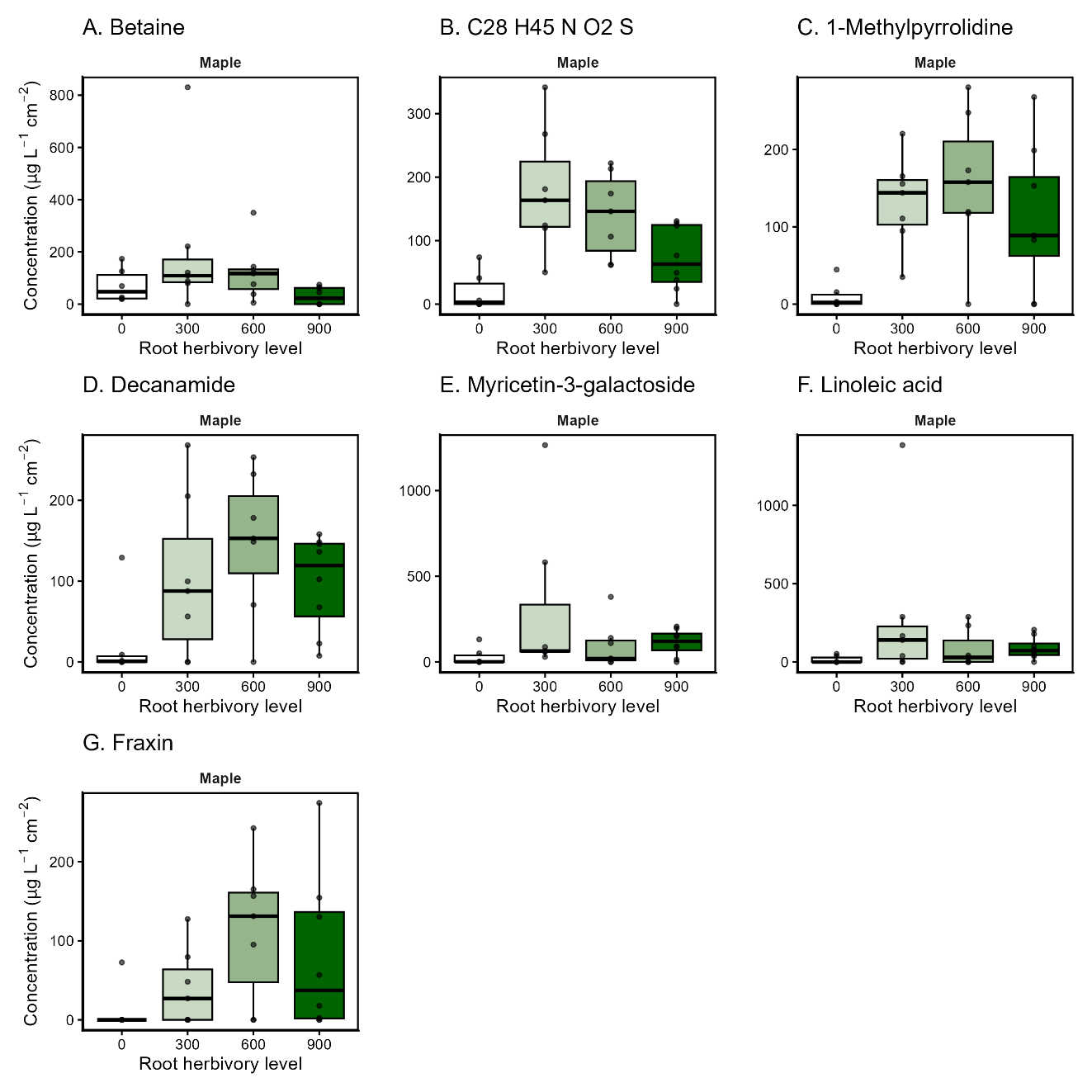

Fig. S10. Concentrations standardized by root area (μg L^-1^ cm^-2^) of individual indicator compounds identified in hickory plots across root herbivory levels (0, 300, 600, and 900 nematodes per microcosm) in positive ionization mode: (A) Betaine, (B) C_28_H_45_NO_2_S, (C) 1-Methylpyrrolidine, (D) Decanamide, (E) Myricetin-3-galactoside, (F) Linoleic acid, and (G) Fraxin (n=56). Boxes represent the IQR, with the centerline indicating the median. Whiskers extend to 1.5 × IQR, and individual points denote raw observations. Color intensity reflects herbivory level, from light (low) to dark (high).

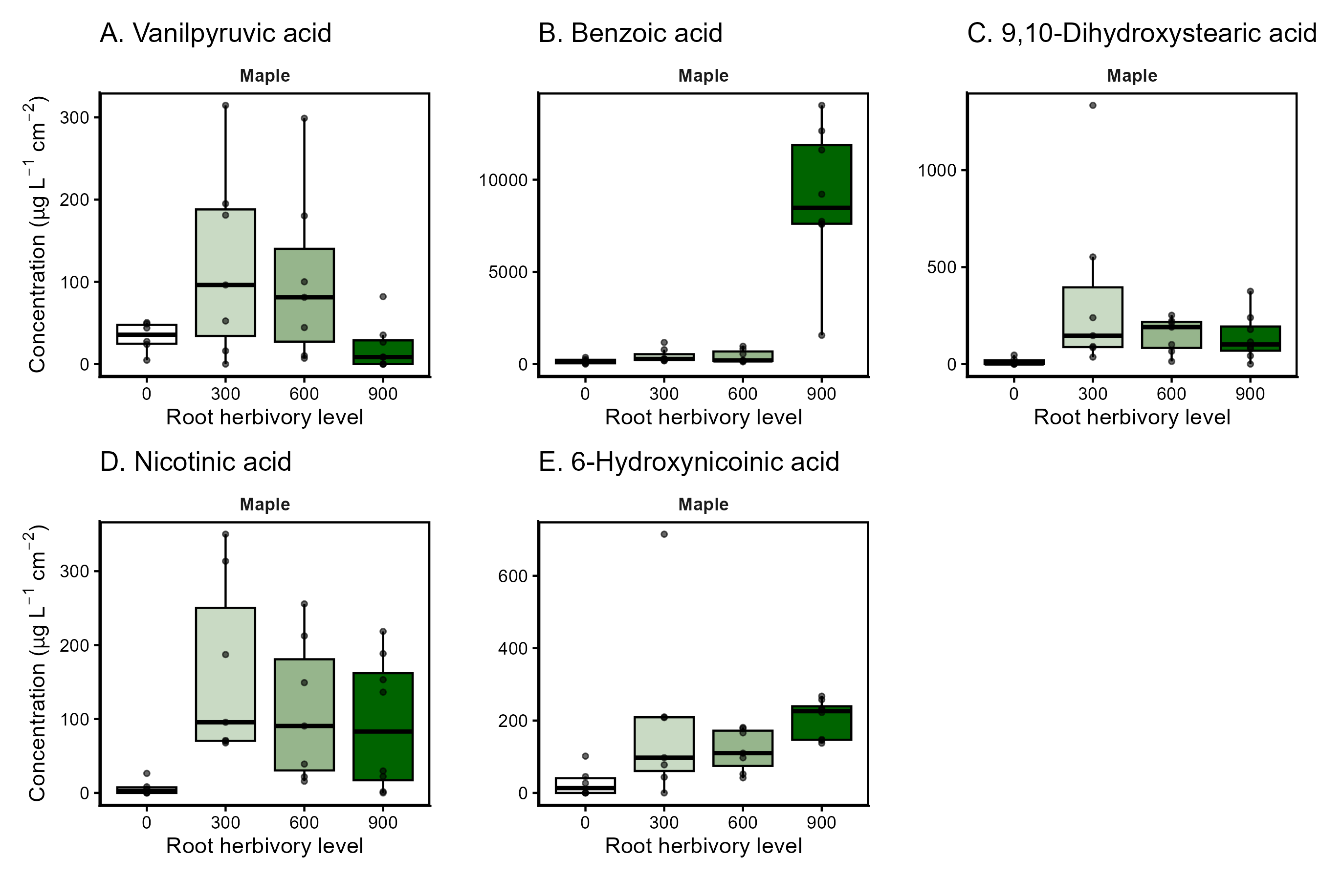

Fig. S11. Concentrations (mg/L) of individual indicator compounds identified in maple plots across root herbivory levels (0, 300, 600, and 900 nematodes per microcosm) in negative ionization mode: (A) Vanilpyruvic acid, (B) Benzoic acid, (C) 9,10-Dihydroxystearic acid, (D) Nicotinic acid, and (E) 6-Hydroxynicotinic acid (n=56). Boxes represent the IQR, with the centerline indicating the median. Whiskers extend to 1.5 × IQR, and individual points denote raw observations. Color intensity reflects herbivory level, from light (low) to dark (high).

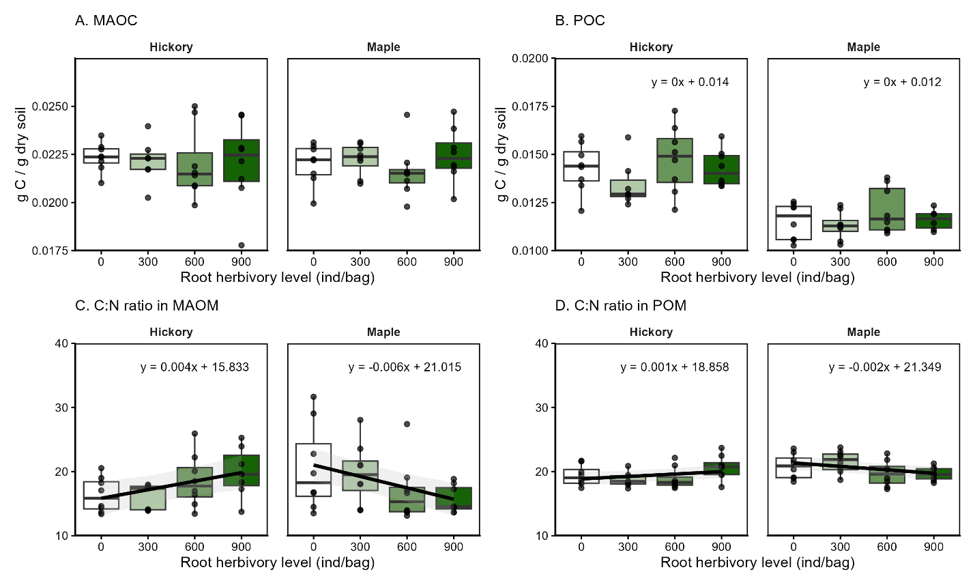
**Fig. S12.** Effects of root herbivory intensity and tree species on soil organic matter fractions and C:N stoichiometry (n=60). Box plots show (A) MAOC (g C/g dry soil), (B) POC (g C/g dry soil), (C) C:N ratio of MAOM, and (D) C:N ratio of POM, across root herbivory levels (0, 300, 600, and 900 nematodes density per microcosm) for hickory (left panels) and maple (right panels). Boxes represent the interquartile range (IQR), with the center line indicating the median. Whiskers extend to 1.5 × IQR, and individual points represent biological replicates. Darker green shading indicates higher herbivory levels. The solid line represents a linear regression fit with 95% confidence intervals (shaded area).

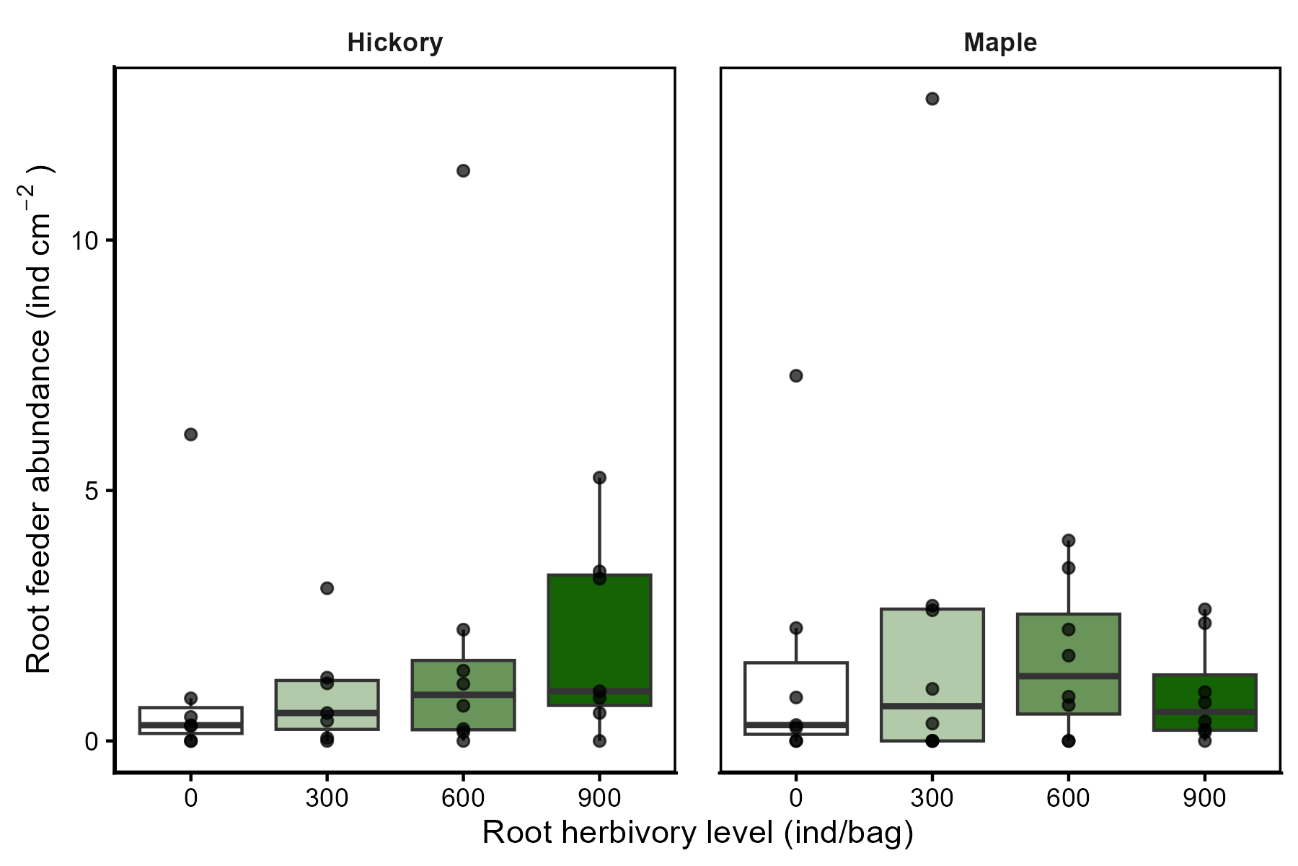
Fig. S13. Post-treatment abundance of root feeding nematodes in roots, standardized by root area (nematode density cm^-2^) across root herbivory levels (0, 300, 600, and 900 nematodes per microcosm) (n=60).

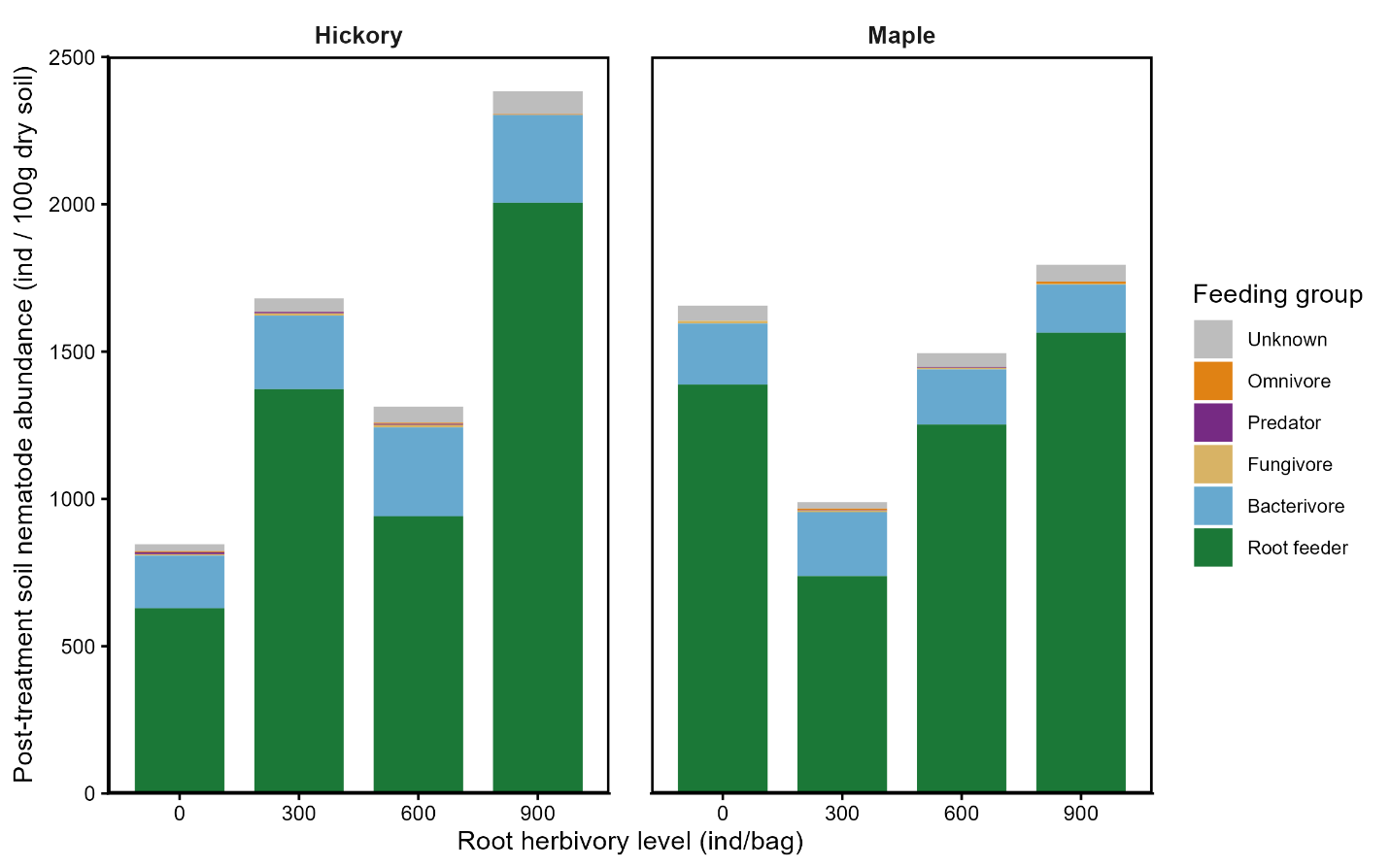

Fig. S14. Mean post-treatment soil nematode trophic group abundance across root herbivory levels (0, 300, 600, and 900 nematodes per microcosm) (nematode density/100g dry soil) (n=60).

Tables

Table S1. Indicator compounds identified in root exudates from hickory and maple trees, their molecular formula and structure, and known biological functions (negative ionization mode). Molecular structure figures are obtained from ChemSpider (14).

| Compound name | Plot | Ionization mode | IndVal  (p-value) | Molecular formula  (average mass) | Biological functions in root exudates |
| --- | --- | --- | --- | --- | --- |
| Vitamin A derivative | Hickory | - | 0.908  (*p*=0.031) |  | (i.e., Apocarotenoids) Root growth regulator, tolerance signal against abiotic and biotic stress (15) |
| Benzoic acid | Maple | - | 0.990  (*p*=0.029) | C_7_H_6_O_2_ (122.123)  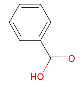 | Salicylic acid signaling, antifungal (16, 17) |
| Nicotinic acid | Maple | - | 0.948  (*p*=0.006) | C_6_H_5_NO_2_ (123.111)  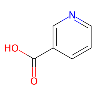 | Nicotinic acid signaling, defense signal against biotic and abiotic stress, increasing glutathione to promote salicylic acid and gibberellin signaling (18) |
| 6-Hydroxynicotinic acid | Maple | - | 0.940  (*p*=0.003) | C_6_H_5_NO_3_ (139.110)  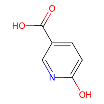 | Nicotinic acid signaling (18) |
| 9,10-Dihydroxystearic acid | Maple | - | 0.949  (*p*=0.005) | C_18_H_36_O_4_ (316.482)  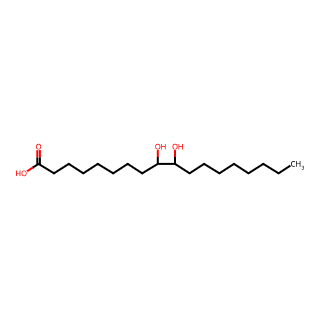 | Hydroxylated fatty acid component of suberin; suberin increases under drought and biotic stress for restructuring the membrane and reinforcing the root cell wall (19) |
| Vanilpyruvic acid | Maple | - | 0.929  (*p*=0.038) | C_10_H_10_O_5_ (210.185)  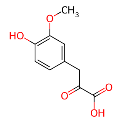 | Unknown root exudate function but potentially related to vanillic acid synthesis (20) |

**Table S2.** Indicator compounds identified in root exudates from hickory and maple trees, their molecular formula and structure, and known biological functions (positive ionization mode). Molecular structure figures are obtained from ChemSpider (14)**.**

| Compound name | Plot | Ionization mode | IndVal  (p-value) | Molecular formula  (average mass) | Biological functions in root exudates |
| --- | --- | --- | --- | --- | --- |
| Jasmonal | Hickory | + | 0.899  (*p*=0.024) | C_14_H_18_O (202.297)  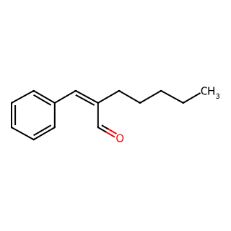 | Jasmonic acid signaling, defense signal against root herbivores, reshaping rhizosphere microbiome (21, 22) |
| 2-Hexenal | Hickory | + | 0.934  (*p*=0.008) | C_6_H_10_O (98.145)  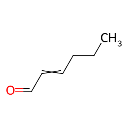 | Jasmonic acid signaling, antibacterial (23–25) |
| Gibberellin | Hickory | + | 0.907  (*p*=0.004) | C_19_H_22_O_6_ (346.379)  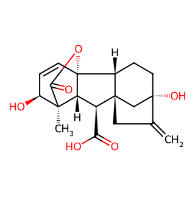 | Jasmonic acid signaling, defense signal against biotic and abiotic stress, restoring root growth (21) |
| Fraxin | Hickory | + | 0.877  (*p*=0.015) | C_16_H_18_O_10_ (370.310)  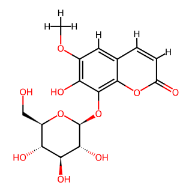 | Iron mobilization under iron-deficient conditions, antimicrobial at wound sites, reshaping rhizosphere microbiome (26) |
|  | Maple | + | 0.810  (*p*=0.046) |  |  |
| Linoleic acid | Maple | + | 0.850  (*p*=0.045) | C_18_H_32_O_2_ (280.452)  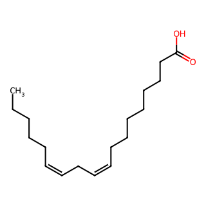 | Jasmonic acid signaling, reshaping rhizosphere microbiome, antimicrobial (27, 28) |
| Betaine | Maple | + | 0.916  (*p*=0.015) | C_5_H_11_NO_2_ (117.148)  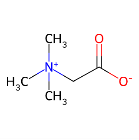 | Defense signal against biotic and abiotic stress, osmotic adjustment under abiotic stress – particularly Glycine betaine (29) |
| Myricetin-3-galactoside | Maple | + | 0.890  (*p*=0.043) | C_21_H_20_O_13_ (480.378)  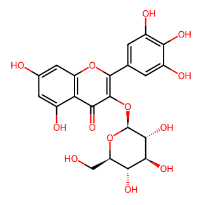 | Reshaping rhizosphere microbiome, antimicrobial (30) |
| 1-Methylpyrrolidine | Maple | + | 0.911  (*p*=0.025) | C_5_H_11_N (85.150)  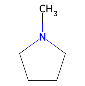 | Unknown root exudate function |
| Decanamide | Maple | + | 0.895  (*p*=0.039) | C_10_H_21_NO (171.284)  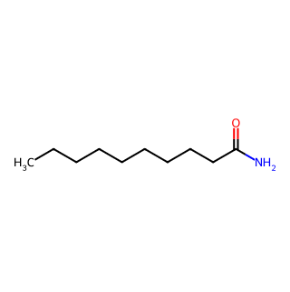 | Unknown root exudate function |
| C_28_H_45_NO_2_S | Maple | + | 0.933  (*p*=0.006) | C_28_H_45_NO_2_S (459.733) | Unknown compound |

**SI References**

1.	A. L. C. Franco, *et al.*, Nematode exclusion and recovery in experimental soil microcosms. *Soil Biology and Biochemistry* **108**, 78–83 (2017).

2.	G. Baermann, Eine einfache Methode zur Auffindung von Ancylostomum (Nematoden) Larven in Erdproben. *Geneeskd Tijdschr Ned Indie* **57**, 131–137 (1917).

3.	J. Flegg, D. Hooper, Extraction of free-living stages from soil. in (1970).

4.	K. C. Ball, *et al.*, Seasonal variation of micro and macro soil invertebrate communities in a temperate forest. *Applied Soil Ecology* **218**, 106729 (2026).

5.	P. A. A. Loof, “The Family Pratylenchidae Thorne, 1949” in *Manual of Agricultural Nematology*, (CRC Press, 1991).

6.	I. A. M. Saeed, A. E. MacGuidwin, D. I. Rouse, Synergism of Pratylenchus penetrans and Verticillium dahliae Manifested by Reduced Gas Exchange in Potato. *Phytopathology®* **87**, 435–439 (1997).

7.	A. L. C. Franco, *et al.*, Root herbivory controls the effects of water availability on the partitioning between above- and below-ground grass biomass. *Functional Ecology* **34**, 2403–2410 (2020).

8.	A. Seethepalli, *et al.*, RhizoVision Explorer: open-source software for root image analysis and measurement standardization. *AoB PLANTS* **13**, plab056 (2021).

9.	J. van Bezooijen, METHODS AND TECHNIQUES FOR NEMATOLOGY. (2006).

10.	J. F. Southey, *Laboratory methods for work with plant and soil nematodes*, 6th Ed. (H.M.S.O, 1986).

11.	B. D. Hafner, O. Pietz, W. L. King, J. B. Scharfetter, T. L. Bauerle, Early developmental shifts in root exudation profiles of five *Zea mays* L. genotypes. *Plant Science* **354**, 112439 (2025).

12.	S. L. Arnold, J. S. Schepers, A Simple Roller-Mill Grinding Procedure for Plant and Soil Samples. *Communications in Soil Science and Plant Analysis* **35**, 537–545 (2004).

13.	L. J. Cihacek, K. A. Jacobson, Effects of Soil Sample Grinding Intensity on Carbon Determination by High‐Temperature Combustion. *Communications in Soil Science and Plant Analysis* **38**, 1733–1739 (2007).

14.	H. E. Pence, A. Williams, ChemSpider: An Online Chemical Information Resource. *J. Chem. Educ.* **87**, 1123–1124 (2010).

15.	A. Felemban, J. Braguy, M. D. Zurbriggen, S. Al-Babili, Apocarotenoids Involved in Plant Development and Stress Response. *Front. Plant Sci.* **10** (2019).

16.	Y. Liu, *et al.*, Three-step biosynthesis of salicylic acid from benzoyl-CoA in plants. *Nature* **645**, 201–207 (2025).

17.	S. Windisch, *et al.*, Role of Benzoic Acid and Lettucenin A in the Defense Response of Lettuce against Soil-Borne Pathogens. *Plants (Basel)* **10**, 2336 (2021).

18.	T. Berglund, A. B. Ohlsson, The nicotinamide hypothesis revisited—plant defense signaling integrating PARP, nicotinamide, nicotinic acid, epigenetics, and glutathione. *FEBS Letters* **599**, 3355–3368 (2025).

19.	J. Braat, *et al.*, The response of Arabidopsis to the apocarotenoid β-cyclocitric acid reveals a role for SIAMESE-RELATED 5 in root development and drought tolerance. *PNAS Nexus* **2**, pgad353 (2023).

20.	T. Yao, *et al.*, Phylogenetic Occurrence of the Phenylpropanoid Pathway and Lignin Biosynthesis in Plants. *Front. Plant Sci.* **12** (2021).

21.	M. Cosme, *et al.*, A fungal endophyte helps plants to tolerate root herbivory through changes in gibberellin and jasmonate signaling. *New Phytologist* **211**, 1065–1076 (2016).

22.	C. Li, *et al.*, Jasmonate Signaling Pathway Modulates Plant Defense, Growth, and Their Trade-Offs. *International Journal of Molecular Sciences* **23**, 3945 (2022).

23.	L. Hu, *et al.*, Herbivory-induced green leaf volatiles increase plant performance through jasmonate-dependent plant–soil feedbacks. *Nat. Plants* **11**, 1001–1017 (2025).

24.	A. Scala, S. Allmann, R. Mirabella, M. A. Haring, R. C. Schuurink, Green Leaf Volatiles: A Plant’s Multifunctional Weapon against Herbivores and Pathogens. *International Journal of Molecular Sciences* **14**, 17781–17811 (2013).

25.	A. Scala, *et al.*, E-2-hexenal promotes susceptibility to Pseudomonas syringae by activating jasmonic acid pathways in Arabidopsis. *Front. Plant Sci.* **4** (2013).

26.	I. A. Stringlis, R. de Jonge, C. M. J. Pieterse, The Age of Coumarins in Plant–Microbe Interactions. *Plant Cell Physiol* **60**, 1405–1419 (2019).

27.	A. Gfeller, L. Dubugnon, R. Liechti, E. E. Farmer, Jasmonate Biochemical Pathway. *Science Signaling* **3**, cm3–cm3 (2010).

28.	T. Wen, M. Zhao, J. Yuan, G. A. Kowalchuk, Q. Shen, Root exudates mediate plant defense against foliar pathogens by recruiting beneficial microbes. *Soil Ecol. Lett.* **3**, 42–51 (2021).

29.	M. G. Annunziata, L. F. Ciarmiello, P. Woodrow, E. Dell’Aversana, P. Carillo, Spatial and Temporal Profile of Glycine Betaine Accumulation in Plants Under Abiotic Stresses. *Front. Plant Sci.* **10** (2019).

30.	G. A. Kumar, *et al.*, Recent advancements in multifaceted roles of flavonoids in plant–rhizomicrobiome interactions. *Front. Plant Sci.* **14** (2024).
